# 3D quantification of zebrafish cerebrovascular architecture by automated image analysis of light sheet fluorescence microscopy datasets

**DOI:** 10.1101/2020.08.06.239905

**Authors:** E. C. Kugler, J. Frost, V. Silva, K. Plant, K. Chhabria, T. J.A. Chico, P. A. Armitage

**Affiliations:** Department of Infection, Immunity and Cardiovascular Disease, University of Sheffield, Medical School, Beech Hill Road, Sheffield, S10 2RX United Kingdom; The Bateson Centre, Firth Court, University of Sheffield, Western Bank, Sheffield, S10 2TN United Kingdom; Insigneo Institute for in silico Medicine, The Pam Liversidge Building, Sheffield, S1 3JD United Kingdom; Hull York Medical School, John Hughlings Jackson Building, University Road, University of York, Heslington, York, YO10 5DD

**Keywords:** angiogenesis, LSFM, registration, quantification, vasculature

## Abstract

Zebrafish transgenic lines and light sheet fluorescence microscopy allow in-depth insights into vascular development *in vivo* and 3D. However, robust quantification of the zebrafish cerebral vasculature in 3D remains a challenge, and would be essential to describe the vascular architecture. Here, we report an image analysis pipeline that allows 3D quantification of the total or regional zebrafish brain vasculature. This is achieved by landmark- or object-based inter-sample registration and extraction of quantitative parameters including vascular volume, surface area, density, branching points, length, radius, and complexity. Application of our analysis pipeline to a range of sixteen genetic or pharmacological manipulations shows that our quantification approach is robust, allows extraction of biologically relevant information, and provides novel insights into vascular biology. To allow dissemination, the code for quantification, a graphical user interface, and workflow documentation are provided. Together, we present the first 3D quantification approach to assess the whole 3D cerebrovascular architecture in zebrafish.

## Introduction

Vascular diseases are the leading cause of death world-wide. Our understanding of the mechanisms that govern vascular development is heavily reliant on preclinical *in vivo* models. Several advantages such as optical clarity of the embryo, coupled with multiple vascular transgenic lines ^1,2^ and advanced microscopy, such as light sheet microscopy (LSFM), makes zebrafish an unrivalled model for dynamic three dimensional (3D) *in vivo* vascular imaging ^3,4^.

Acquisition of detailed imaging datasets of zebrafish vasculature has advanced considerably, however objective reliable comparison and quantification of these datasets is still limited, with most studies relying on subjective visual assessment rather than objective quantification. The lack of robust automated approaches to quantify vascular anatomy in 3D is a substantial limitation, reducing analytical throughputs and preventing detection of subtle phenotypes.

Development of an automated 3D vascular quantification pipeline requires many obstacles to be overcome. LSFM datasets are often terabytes in size, rendering data handling, processing, and analysis computationally demanding. Analysis of preclinical imaging data also often lacks an understanding of data properties such as noise, artefacts and motion. A further challenge is that, typically, only 3-15% of voxels in these datasets are vascular, limiting the amount of information available to drive analysis approaches ^5–7^. Together, these issues have led most quantitative studies to analyse 2D projections, failing to capture the complexity of the 3D vasculature, and is confounded by overlying vessels. 3D quantification would clearly be accurate, but is technically and computationally more challenging.

Very few previous studies have attempted to quantitatively characterise the zebrafish cranial vasculature. Tam *et al.* ^8^ and Chen *et al.* ^9^ focussed on characterising small vascular sub-regions in confocal microscopy images using commercial software. We previously presented a 3D enhancement and segmentation approach for the cranial vasculature in zebrafish transgenic reporter lines ^10,11^. Daetwyler *et al.* ^12^ presented a method to segment the zebrafish cerebral vasculature using machine learning (dual ResUNet), but did not include further quantification.

Most recently we presented a validation of our segmentation method ^13,14^.

We here present the first easily applicable 3D image analysis pipeline for zebrafish vasculature. We apply this to the cerebral vasculature with **(a)** registration that allows examination of vascular patterning similarity/variability between individuals or groups, and **(b)** perform global and regional quantification of eight vascular parameters (*i.e. similarity, volume, surface area, density, branching points, length, radius, and complexity*) (**Fig. 1**).

**Figure 1:**
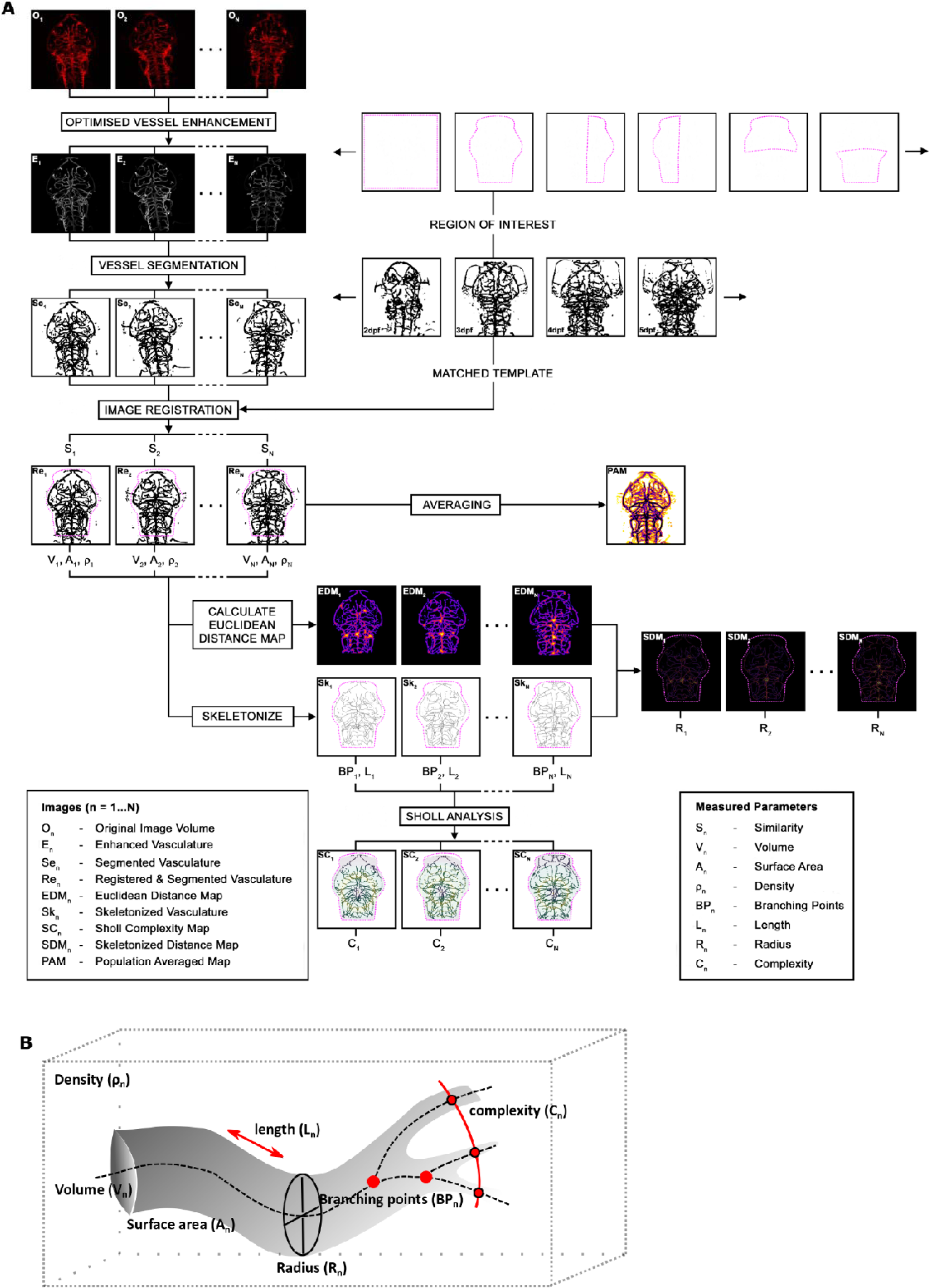
Workflow for 3D image registration and quantification. **A** The original images (O_n_) were motion-corrected, enhanced (En), and segmented (S_en_). Registration to template embryo brought all embryos into one spatial coordinate system (Re_n_) allowing the quantification of similarity (S_n_), and followed by averaging (PAM_n_). Following ROI extraction, volume (V_n_), surface voxel (A_n_), and density (ρ_n_) were quantified. EDMs (EDM_n_) were combined with vascular skeletons (Sk_n_) for radius quantification (SDM_n;_ R_n_), while skeletons were used to quantify network length (L_n_), branching points (BP_n_), and complexity (C_n_). **B** Schematic of the parameters extracted.

To demonstrate the utility and throughput of our approach, we quantified **(a)** embryonic vascular development from 2-to-5dpf, expecting changes due to development, **(b)** the impact of lack of blood flow ^15^, expecting zebrafish without blood flow to display an impaired vascular architecture, **(c)** the effect of sixteen genetic manipulations, which were expected to impact angiogenesis (via previously described morpholinos targeting *jagged-1a* ^16^*, jagged-1b* ^16^*, dll4* ^16^*, notch1b* ^17^*, ccbe1*^18^), **(d)** pharmacological manipulations of components associated with angiogenesis, vasoconstriction, and endothelial cell (EC) architecture (vascular endothelial growth factor (VEGF) inhibition ^19^, Notch inhibition ^20^ nitric oxide synthase (NOS) inhibition ^21^, Wnt inhibition ^22^, Wnt activation ^23^, F-actin polymerization inhibition ^24^, Myosin II inhibition ^25^, osmotic pressure increase, membrane rigidity decrease) and **(e)** examine regional topology of the mid-versus hindbrain and left-right vascular symmetry.

To allow dissemination and wider application of our method, we have implemented it as a workflow in the open-source image analysis software Fiji ^26^, containing a custom graphical user interface (GUI), accompanied by comprehensive workflow documentation.

In summary, we describe the first comprehensive 3D quantification approach for zebrafish cerebrovascular characterisation that would have potential widespread applications and will aid discovery of novel insights into vascular development.

## Results

### Vascular inter-sample registration allows identification of similar vessel patterns between embryos

We first applied our optimised segmentation approach to the transgenic *Tg(kdrl:HRAS-mCherry)^s916^* ^27^ (Sato enhancement and Otsu thresholding ^11,28,29^).

To identify landmarks for registration, we performed manual measurements to gain insights into the extent of vascular similarity (**Fig. S1**). This showed that vascular growth was highly comparable between embryos and that a landmark-based registration could be applicable for registration. This visual inspection allowed identification of eleven anatomical landmarks for registration, shown in **Fig. 2A-C**.

**Figure 2:**
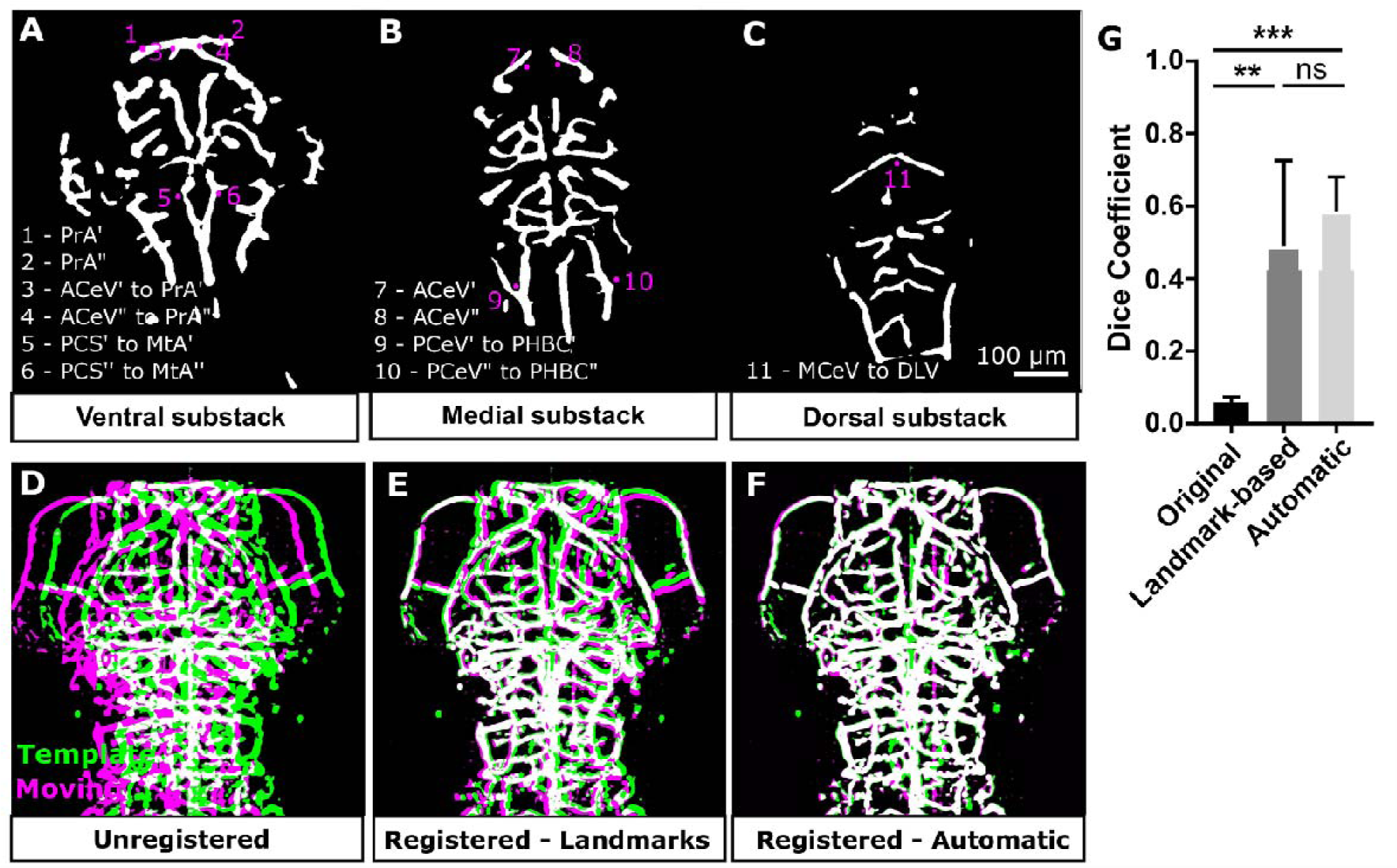
Testing and validation of rigid landmark- or object-based inter-sample registration. **(A-C)** Anatomical landmarks chosen for landmark-based rigid registration were as follows: (1) left prosencephalic artery (PrA’), (2) right PrA (PrA’’), (3) junction point of left PrA and anterior cerebral vein (ACeV’), (4) junction point of right PrA’’ and ACeV’’, (5) junction of left posterior communicating segment (PCS’) to metencephalic artery (MtA’), (6) junction of right PCS’’ to MtA’’, (7) highest curvature point of ACeV’, (8) highest curvature point of ACeV’’, (9) left posterior junction point of posterior cerebral vein (PCeV’) and primordial hindbrain channel (PHBC’), (10) left posterior junction point pf PCeV’’ and PHBC’’, (11) junction point of middle cerebral vein (MCeV) and dorsal longitudinal vein (DLV). Positions are indicated in representative image. **D** Same sample from two acquisitions overlaid following segmentation (green - template; magenta - moving image; white - overlap). **E** Samples after rigid anatomical landmark-based registration. **F** Samples after rigid automatic registration. **G** Dice coefficient was statistically significantly increased after landmark-based (p 0.0017) and automatic registration (p 0.0003; n=5 embryos; 3 experimental repeats; One-Way ANOVA).

We next used both, manual rigid landmark-based registration and an automatic rigid registration approach ^30,31^ to bring segmented images into a single spatial coordinate system, allowing us to compare two methods against each other. To validate their applicability both, manual landmark-based and automatic registration, were applied to multiple images of the same sample that was manually displaced between consecutive image acquisitions. Both registration approaches significantly increased similarity, quantified by Dice coefficient (landmark-based p 0.0017 and automatic p 0.0003; One-Way ANOVA; **Fig. 2D-G**), with a lower coefficient of variation following automatic (16.23%) than landmark-based registration (48.79%).

To compare registration outcomes when applied to different embryos, both registration methods were applied to register six embryos from 2-to-5dpf (by registering five individuals to another single target embryo) (**Fig. S2**), allowing the identification of regions of vascular similarity across embryos (**Fig. 3A-C; Video S1-S4**). Quantification of Dice score showed higher similarity after automatic registration (**Fig. 3D**; 2dpf manual p>0.9999, automatic p 0.2154; 3dpf manual p>0.9999, automatic p<0.0001; 4dpf manual p>0.9999, automatic p 0.3962; 5dpf manual p>0.9999, automatic p>0.9999) and a lower coefficient of variation (CoV) was found after automatic registration (2dpf 38.18%, 3dpf 26.21%, 4dpf 29.23%, 5dpf 17.93%) than manual registration (2dpf 84.02%, 3dpf 81.20% 4dpf 26.21%, 5dpf 18.39%). Moreover, averaging allowed examination of regions of similarity and variability (**Fig. S2D-G**), showing that the main cerebral vessels, such as the basilar artery (BA), have a high degree of anatomical similarity between fish.

**Figure 3:**
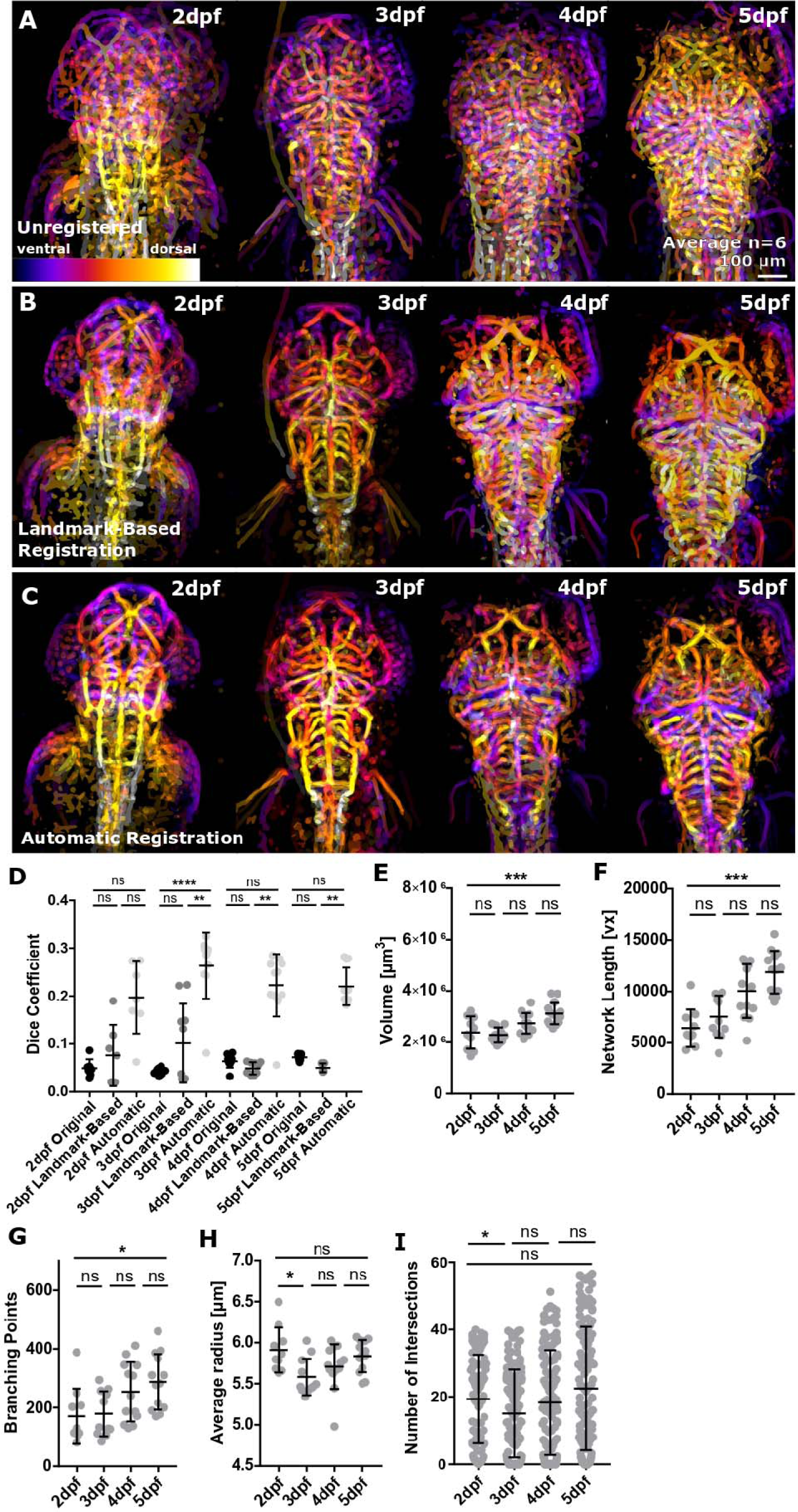
Inter-sample registration from 2-to-5dpf enables the study of regions of similarity and variability. **A** Depth-coded MIP showing regions of overlap (purple - ventral, white - dorsal) of six fish before registration from 2-5dpf. **B** Depth-coded MIP showing regions of overlap (purple - ventral, white - dorsal) of six fish after manual registration from 2-5dpf. **C** Depth-coded MIP showing regions of overlap (purple - ventral, white - dorsal) of six fish after automatic registration from 2-5dpf. **D** Dice coefficient between template and moving image was increased after application of rigid registration using both anatomical landmark-based and automatic rigid registration from 2-to-5dpf (2dpf n=7, 3dpf n=10, 4dpf n=10, 5dpf n=10; 2 experimental repeats; Kruskal-Wallis test; ns p>0.5, *p 0.05-0.01, **p 0.01-0.001, ***p 0.001-0.0001, ****p<0.0001). **E** Vascular volume was statistically significantly increased from 2-to-5dpf (p 0.0008; 2dpf n=10, 3dpf n=12, 4dpf n=13, 5dpf n =15; One-Way ANOVA). **F** Vascular network length was statistically significantly increased from 2-to-5dpf (p 0.0001; 2dpf n=10, 3dpf n=12, 4dpf n=13, 5dpf n =15; Kruskal-Wallis test). **G** Branching points were statistically significantly increased from 2-to-5dpf (p 0.0082; Kruskal-Wallis test). **H** Average vessel radius was not statistically significantly changed from 2-to-5dpf (p>0.9999; Kruskal-Wallis test). **I** Sholl analysis was conducted to assess vascular complexity, showing no significant increase from 2-to-5dpf (2-3dpf p 0.0340, 3-4dpf p 0.6825, 4-5dpf p 0.2000; 2-5dpf n=5; Kruskal-Wallis test).

This demonstrated that landmark- and object-based rigid inter-sample registration can be used to bring the vasculature of different embryos into one spatial coordinate system, and is applicable from 2-to-5dpf. Moreover, automatic registration performs better than manual landmark-based registration. Thus, a common coordinate system allows for consistent automated placement of regions of interest, which improves speed and reproducibility of quantification.

### Establishment of 3D vascular parameter quantification

We established a workflow in the open-source image analysis framework, Fiji ^26^ to quantify **(i)** similarity (S_n_): volume of overlap quantified by Dice Coefficient, **(ii)** vascular volume (V_n_): vascular voxels derived following voxel-classification as vascular and non-vascular using segmentation, **(iii)** surface area (S_n_), **(iv)** density (ρ_n_): ratio of vascular volume to total volume of interest, **(v)** network length (L_n_): number of centreline voxels, derived by 3D-thinning of the segmented vasculature, **(vi)** branching points (BP_n_): points where vessels splits up into daughter branches, **(vii)** average radius (R_n_): vascular thickness given by the distance from local centreline (or vessel radius midpoint) to corresponding vessel walls, and **(viii)** complexity (C_n_): number of concentric shell intersections from centre point using Sholl analysis, where the subscript *n* represents a measurement in the n^th^ fish of the group under study (**Fig. 1**).

The established quantification pipeline was applied to data from 2-to-5dpf which were expected to show changes in vascular topology over time with increasing volume, length, branching points, and complexity. Quantification showed that vascular volume (**Fig. 3E**; p 0.0008; One-Way ANOVA), network length (**Fig. 3F**; p 0.0001; Kruskal-Wallis test), and branching points (**Fig. 3G**; p 0.0082; Kruskal-Wallis test), but not average radius (**Fig. 3H**; p>0.9999; Kruskal-Wallis test) and complexity (**Fig. 3I**; p>0.9999; Kruskal-Wallis test), significantly increased from 2-to-5dpf. This suggested that our quantification approach was applicable for extraction of biologically relevant vascular parameters.

### Quantification of the effect of absence of blood flow on vascular anatomy

To establish whether our workflow successfully extracts biologically relevant vascular differences in embryos of the same age, we applied it to analyse 3dpf zebrafish with or without blood flow (achieved by *tnnt2a* morpholino antisense knockdown; **Fig. 4A**). Quantification showed that Dice coefficient (**Fig. 4B**), vascular volume (**Fig. 4C**), vascular surface area (**Fig. 4D**), branching points (**Fig. 4F**), network length (**Fig. 4G**), average vessel radius (**Fig. 4H**) and vascular complexity were statistically significantly decreased in *tnnt2a* morphants (**Fig. 4I**). The only parameter not statistically significantly changed was vascular density (**Fig. 4E**). This suggests that our quantification detects biological differences of relevant effect size in embryos of the same age-group with commonly used group sizes.

**Figure 4:**
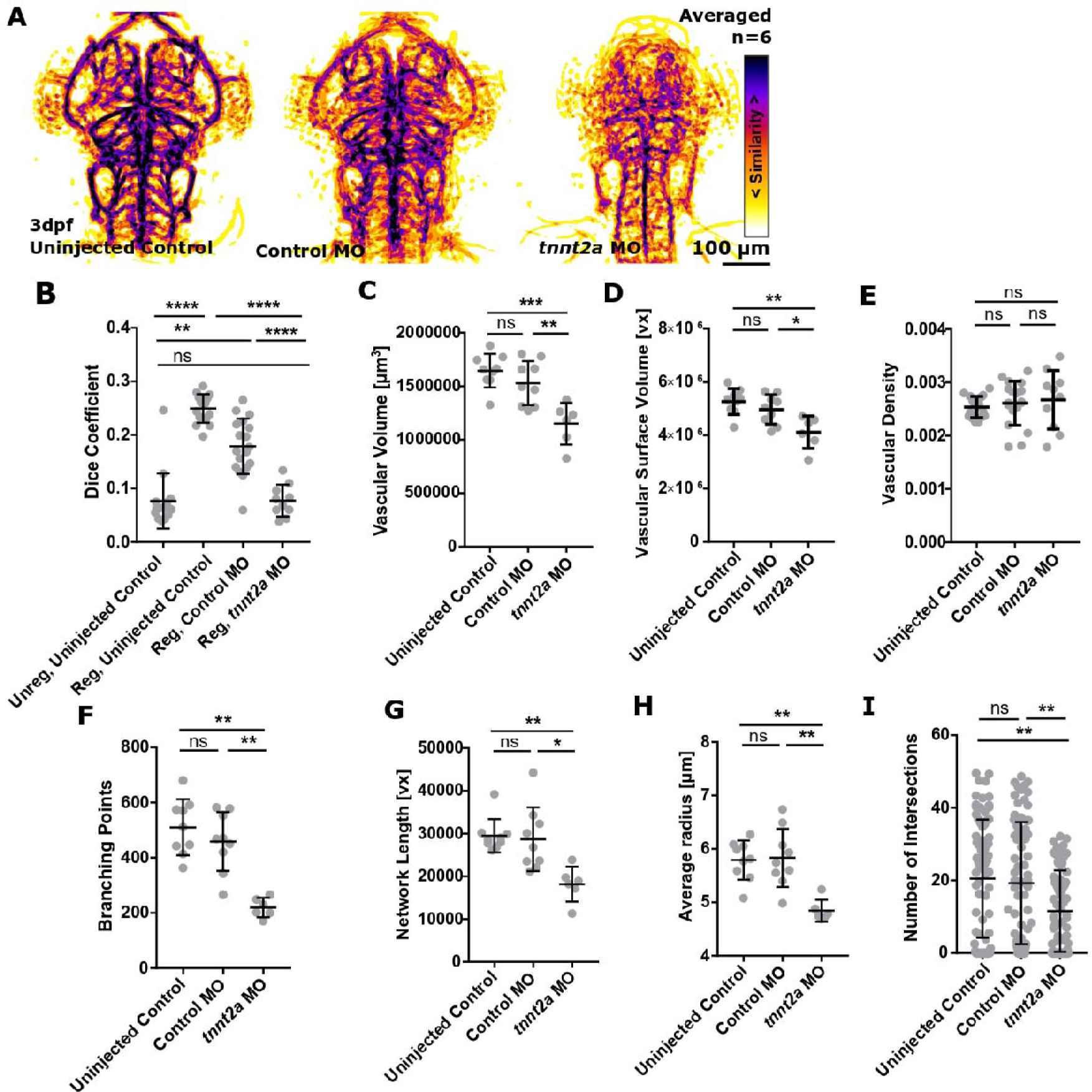
The impact of blood flow loss on vascular topology. **A** MIPs of averaged data of uninjected controls, control MO, and *tnnt2a* MO following segmentation and registration. **B** A statistically significant decrease in Dice coefficient was found when comparing registered control MO to *tnnt2a* MO (p<0.0001; uninjected control=15, control MO=18, *tnnt2a* MO=10; Kruskal-Wallis test). **C** Vascular volume was statistically significantly decreased in *tnnt2a* MO (uninjected control p<0.0001, control MO p<0.0001; uninjected control=9, control MO=9, *tnnt2a* MO=6; One-Way ANOVA). **D** Vascular surface was statistically significantly decreased in *tnnt2a* MO (uninjected control p<0.0001, control MO p 0.0007; Kruskal-Wallis test). **E** Vascular density was not statistically significantly changed in *tnnt2a* MO (uninjected control p 0.6514, control MO p 0.9082; One-Way ANOVA). **F** Branching points were statistically significantly decreased in *tnnt2a* MO (uninjected control p 0.0019, control MO p 0.0092; Kruskal-Wallis test). **G** Vascular network length was statistically significantly decreased in *tnnt2a* MO (uninjected control p 0.0033, control MO p 0.0209; Kruskal-Wallis test). **H** Average vessel radius was statistically significantly decreased in *tnnt2a* MO (uninjected control p 0.0050, control MO p 0.0067; Kruskal-Wallis test). **I** Vascular complexity was statistically significantly decreased in *tnnt2a* MO (uninjected control p 0.0050, control MO p 0.0067; Kruskal-Wallis test).

### 3D quantification allows novel insights into the role of proteins, signalling pathways, and chemical modulators

Our analysis workflow was then applied to existing datasets to quantify the effect of a range of morpholino-induced gene knock-down and pharmacological manipulation (**Fig. 5A**; **Table S1**).

**Figure 5:**
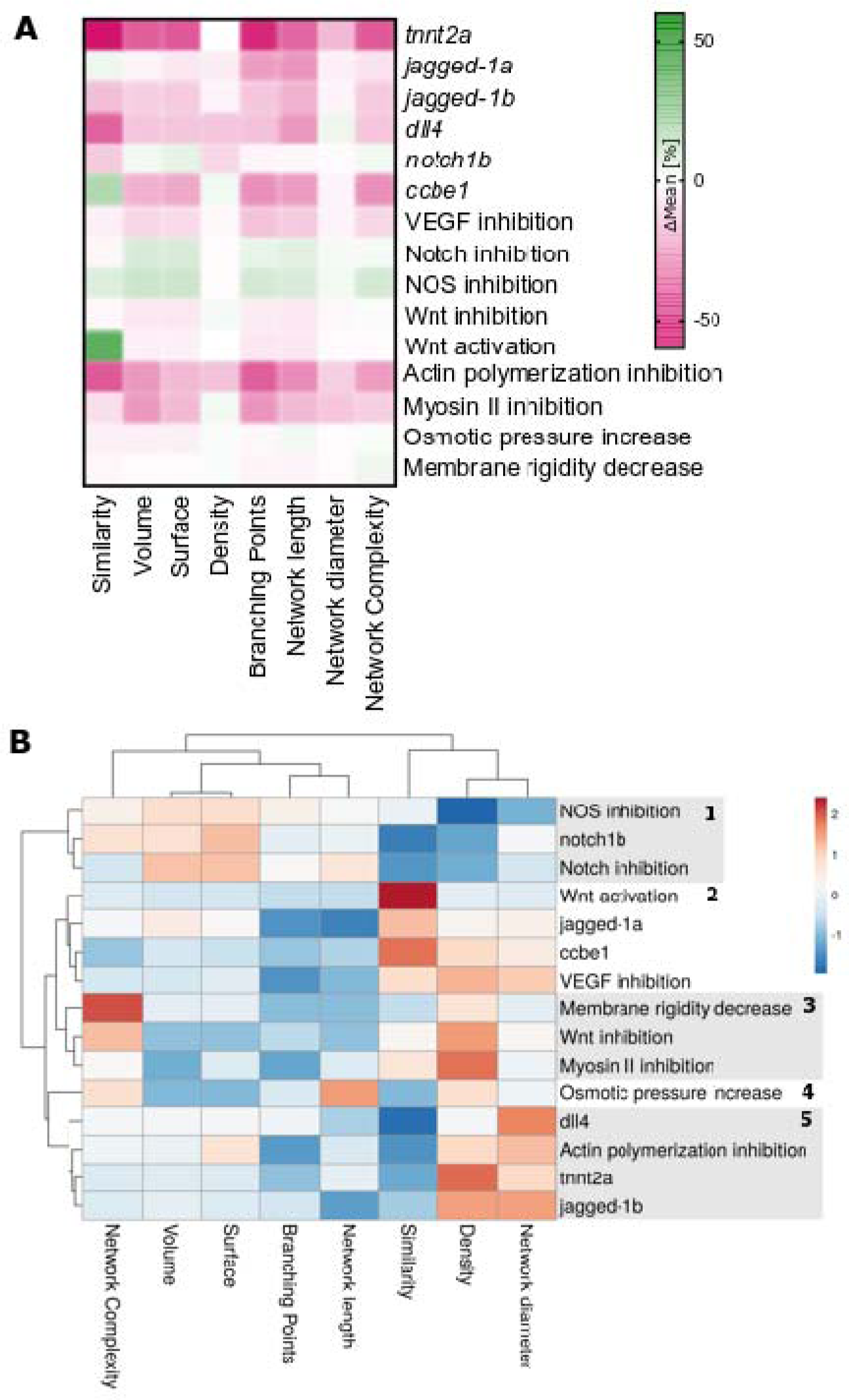
Quantification of vascular parameters and cluster analysis. **(A)** Percentage differences in mean values were quantified for the measured parameters (control MO to MO, and untreated to treated). **(B)** Cluster analysis identified four main clusters, including (1) inhibitors of anti-angiogenic factors, (2) inhibitors of angiogenic factors, (3) factors inducing cellular changes, (4) osmotic pressure, and (5) factors with severe angiogenic defects.

Jagged1 was previously shown to be a proangiogenic regulator in mice, but so far no study examined the distinctive roles of the zebrafish orthologues *jagged-1a* or *jagged-1b* in zebrafish angiogenesis. Our data show that loss of *jagged-1a* leads to decreased network length and branching points (**Fig. S3**), while loss of *jagged-1b* results in reduced vascular volume, surface area, and network length (**Fig. S4**).

We next examined the impact of *dll4* loss, which was previously shown to result in hyper-vascularization in the zebrafish trunk ^32^ and mouse retina ^33^. Our quantification shows that the opposite is the case in the zebrafish brain vasculature with a decrease of all vascular parameters, except radius, and complexity (**Fig. S5**).

Loss of *notch1b* was previously shown to not significantly impact vascular topology in the zebrafish trunk ^17^, which is reproduced by our data (**Fig. S6**).

Previous work showed that *ccbe1* is required for lymphangiogenesis and venous differentiation ^34^, and loss of *ccbe1* impacts murine coronary artery ^35^. Our quantification of vascular topology upon *ccbe1* loss shows a significant reduction of volume, surface area, branching points, and network length (**Fig. S7**).

We next examined the impact of chemically inhibiting VEGF signalling, as VEGF signalling is considered to be a pro-angiogenic factor ^36–38^. Our quantification revealed that already after two hours of VEGF-receptor (VEGFR) inhibition almost all measured vascular parameters are statistically significantly reduced (AV951, **Fig. S8**).

Contrarily, Notch signalling is considered to be anti-angiogenic and loss of Notch to result in hyper-vascularisation ^39,40^. Our data show an increase of almost all measured vascular parameters, but only a statistically significant increase in vascular volume and surface area upon Notch inhibition (DAPT, **Fig. S9**).

Following, we were interested in the impact of NOS inhibition as this was previously shown to result in vasoconstriction ^41^. However, our data showed no statistically significant change in any measured parameters (L-NAME, **Fig. S10**).

We next studied the impact of Wnt inhibition (XAV939) and Wnt activation (GSK3 inhibitor), as Wnt signalling is crucial for cerebrovascular development and blood-brain-barrier development ^42–44^. Our data showed that short-term (4h) Wnt inhibition or Wnt activation does not lead to a statistically significant change of any of the quantified vascular parameters (**Fig. S11, S12**).

We then examined the impact of inhibiting F-actin polymerization and Myosin II cytoskeletal components, which are pivotal for EC stability and also indirectly impact the vascular lumen via vascular smooth muscle cell (vSMC) contractile machinery ^45^. When quantifying vascular parameters upon inhibition of F-actin polymerization (Latrunculin B) and Myosin II (Blebbistatin) a statistically significant reduction of almost all parameters was found (**Fig. S13, S14**).

Next, we studied the impact of increased osmotic pressure using glucose which is thought to result in cellular swelling, potentially impacting angiogenesis ^46,47^, with a previous study in zebrafish showing that only exposure over 96h impacted vascular topology in tectal vessels ^48^. Our data confirm 24h glucose exposure does not impact cerebrovascular topology significantly (**Fig. S15**).

Lastly, we examined the impact of decreased membrane rigidity (DMSO) on vascular topology as it increases membrane permeability ^49^, results in transient osmotic absorption in vessels ^50^ and increases membrane permeability ^49^, which could potentially change at least vessel radius. However, our data suggest that a 24h exposure does not lead to statistically significant changes of vascular topology (**Fig. S16**).

To further examine the impact of the examined morpholinos and chemicals on vascular topology, we quantified percentage differences in mean values of all of the measured quantitative parameters (**Fig. 5A**; **Table S1**; control MO to MO, and untreated to treated; see **Videos S5-S19**), whereas percentage differences were used to allow comparability between groups.

Cluster analysis showed that five main clusters existed, which can be considered as **(1)** inhibitors of anti-angiogenic factors (NOS inhibition, *notch1b* MO, Notch inhibition), **(2)** inhibitors of angiogenic factors (Wnt activation, *jagged-1a* MO, *ccbe1* MO, VEGF inhibition), **(3)** factors inducing cellular changes (membrane rigidity increase, Wnt inhibition, Myosin II inhibition), **(4)** osmotic pressure, and **(5)** factors with severe angiogenic defects (*dll4* MO, actin polymerization inhibition, *tnnt2a* MO, *jagged-1b* MO) (**Fig. 5B**).

Our data showed that meaningful parameters can be extracted to quantify and describe vascular topology. Together, our work presents by far the most comprehensive study quantifying vascular changes upon genetic loss of function and chemical inhibition/activation.

### Comparing regional similarity and left-right symmetry

Next, we examined regional vascular topology. Our initial manual measurements of local vascular growth had shown that growth was comparable between fish (**Fig. S1**), thus embryos could be compared to each other on a local scale.

We therefore quantified vascular parameters from automatically prescribed midbrain and hindbrain regions at 3dpf (**Fig. 6A**). As this has never been done before, it was unclear whether vessel radius, network length, branching points, or volume would be similar or dissimilar when compared in the same zebrafish. We found vascular network length (p 0.6168; **Fig. 6B**), branch points (p 0.0717; **Fig. 6D**), and vascular volume (p 0.0717; **Fig. 6E**) to be not significantly different, while the average vessel radius was significantly higher in the hindbrain (p 0.0008; **Fig. 6C**). Comparing vessel radii further, the BA was found to be the main contributing factor for the increased average vessel radius in the hindbrain (**Fig. S18**).

**Figure 6:**
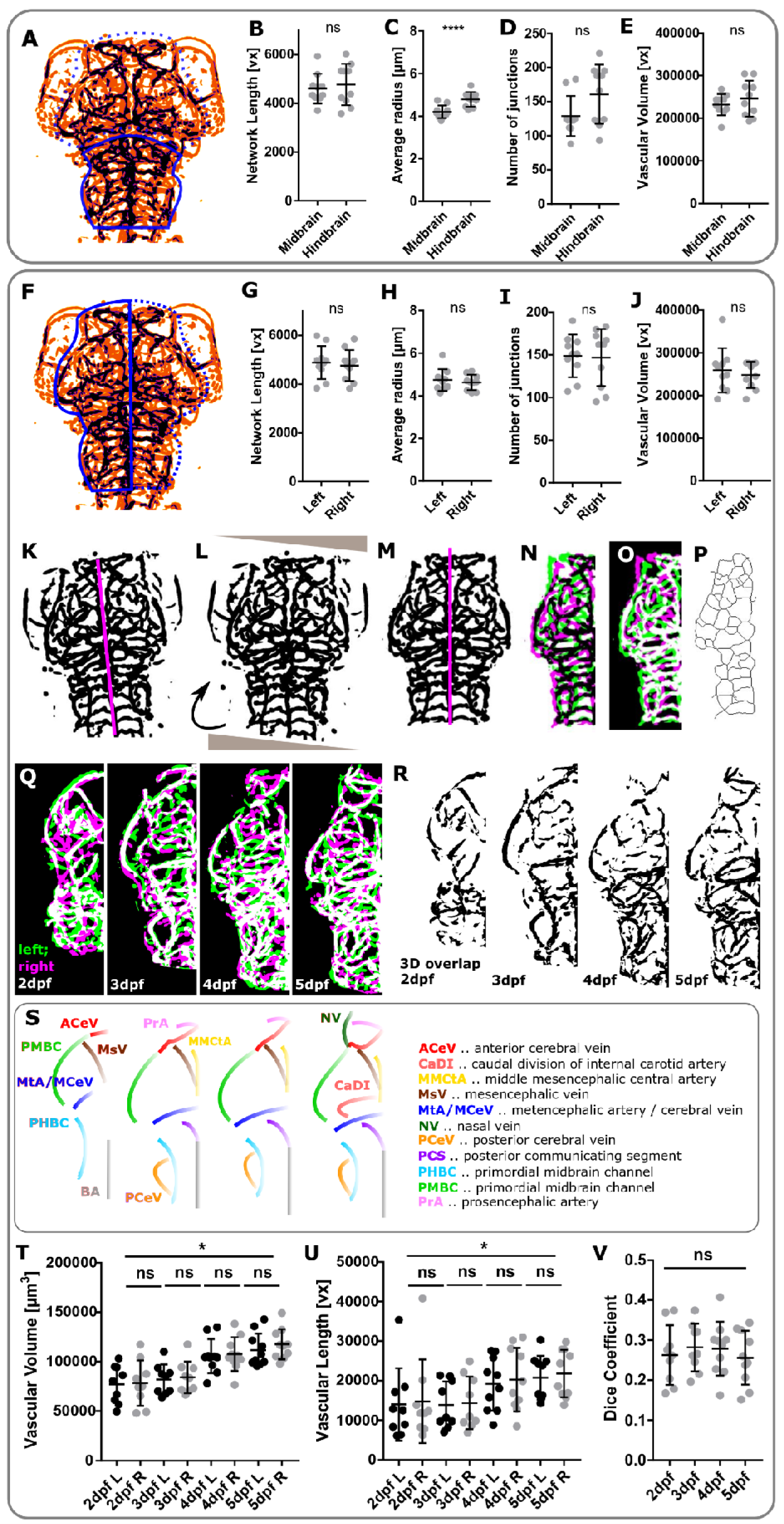
Comparing regional similarity and left-right symmetry. **A** ROI selection allows comparison of vascular parameters in the midbrain (dashed line) and hindbrain (solid line). **B** Comparing the midbrain to the hindbrain, network length was not statistically significantly different (p 0.6168, n=10, unpaired Student’s t-test). **C** The average vessel radius was statistically significantly higher in the hindbrain (p 0.0008, n=10, unpaired Student’s t-test). **D** The number of junctions was not statistically significantly different (p 0.0717, n=10, unpaired Student’s t-test). **E** Vascular volume was not statistically significantly different (p 0.3698, n=10, unpaired Student’s t-test). **F** ROI selection allows comparison of vascular parameters in the right (dashed line) and left (solid line) brain hemisphere. **G** Comparing the left-to-right vasculature, network length was not statistically significantly different (p 0.6830, n=10, unpaired Student’s t-test). **H** The average vessel radius was not statistically significantly different (p 0.5665, n=10, unpaired Student’s t-test). **I** The number of junctions was not statistically significantly different (p 0.8816, n=10, unpaired Student’s t-test). **J** Vascular volume was not statistically significantly different (p 0.5730, n=10, unpaired Student’s t-test). **K** Manual selection of left-right body axis (magenta) in segmented template fish used for image rotation and left-right separation in registered segmented images. **L** Image rotation was performed to align sample anterior-posterior axis with image x-axis. **M** Right vascular volume was mirrored and vascular volume quantified for the left and right vasculature. **N,O** Similarity measures were extracted to compare left and mirrored right vasculature. **P** Left and right vascular network length were quantified after skeletonization to extract vascular centrelines (representative images). **Q** MIPs of left (green) and right (magenta) vasculature, showing regions of similarity (white). **R** Representative mircographs of regions of left-right vascular overlap from one example fish. **S** Schematics showing left-right symmetric vessels from 2-to-5dpf. **T** Cranial vascular volume was found to statistically significantly increase from 2-to- 5dpf (p<0.0001), while no statistically significant difference was found between the vascular volume of the left (L) and right (R) brain hemisphere (L-R 2dpf p<0.9999, 3dpf p>0.9999, 4dpf p>0.9999, 5dpf p 0.9946; embryos n 2dpf=9, 3dpf=9, 4dpf=9, 5dpf=10; 2 experimental repeats; One-way ANOVA). **U** Cranial vascular network length was found to statistically significantly increase from 2-to-5dpf (p 0.0194), while no statistically significant L-R difference was found (L-R 2dpf p<0.9999, 3dpf p>0.9999, 4dpf p>0.9999, 5dpf p>0.9999; samples as above; Kruskal-Wallis test). **V** Dice coefficient was not statistically significantly changed between the left and right vasculature from 2-to-5dpf (p 0.8058; One-way ANOVA; embryos n 2dpf=9, 3dpf=9, 4dpf=10, 5dpf=10; data from 2 experimental repeats).

Comparing the left-to-right vasculature at 3dpf (**Fig. 6F**), no statistically significant difference in network length (p 0.6830; **Fig. 6G**), average vessel radius (p 0.5665; **Fig. 6H**), number of junctions (p 0.8816; **Fig. 6I**), or vascular volume (p 0.5730; **Fig. 6J**) was found.

To examine left-right vascular symmetry in more detail and over time, we then developed an image analysis workflow to assess topological similarity of the left and right vasculature automatically (**Fig. 6K-P**). This was achieved by mirroring the vasculature from the right brain hemisphere to the left by horizontal transformation (**Video S20**) and extracting vessel overlap by voxel majority decisions (**Fig. 6Q,R**). This suggested that peripheral/perineural vessels show a strong degree of left-right symmetry from 2-to-5dpf (**Fig. 6S**). Quantification of vascular volume (**Fig. 6T**) and network length (**Fig. 6U**), showed no statistically significant difference from 2-to-5dpf, although the Dice coefficient between the left and right vasculature was low (**Fig. 6V**).

Thus, our quantification approach allows for novel regional insights of vascular topology, showing that the average vessel radius is higher in hindbrain vessels than midbrain vessels, but that the vasculature is highly symmetric from 2-to-5dpf.

## Discussion

In this work, we present the first biomedical image analysis workflow that allows 3D zebrafish cerebrovascular quantification, including **(a)** inter-sample registration, **(b)** quantification of parameters (vascular volume, surface area, density, network length, branching points, radii, and network complexity), and **(c)** quantification of regional vascular parameters comparing midbrain-to-hindbrain and left-to-right vascular topology, using the freely available image analysis software framework Fiji.

Quantification of Dice scores following registration resulted in low Dice scores for both registration methods, but no reference was available to compare our results to. We anticipate that data sparsity of the segmented vasculature is one contributing factor for these low Dice scores, as small shifts between vessels caused either by inter-sample variation or residual registration errors will affect the overlap more drastically than it would be the case for solid structures such as the brain. Future work is needed to examine Dice coefficients on vessel-segment levels, study voxel-wise comparisons, explore alternative similarity metrics more suited to a sparse vascular network, and develop registration methods that utilise higher order transformations.

Our registration approach allows for the examination of regional similarity as well as facilitating development of a vascular growth atlas. Future work could use double transgenics to allow other information, such as expression patterns, to be co-registered to the vascular growth atlas, as has been performed for a zebrafish brain atlas ^51,52^.

Examining the loss of *jagged-1a* or *jagged-1b*, we show for the first time that loss of either results in mild cerebrovascular defects. As *jagged-1a* and *jagged-1b* are paralogues found in zebrafish due to partial genome duplication ^53^, we anticipate that a loss of both at the same time might result in a stronger effect.

Finding that *dll4* knock-down results in a reduction of the cerebral vasculature is in contrast to previous work in the zebrafish trunk ^32^. This suggests that *dll4* might have cerebrovascular-specific roles which were previously overlooked and require future investigation.

Our finding that loss of *notch1b* does not significantly impact vascular topology suggests that *notch1b* plays a conserved role in the zebrafish head and trunk vasculature ^17^.

Having found that loss of *ccbe1* results in a decrease of vascular volume, surface area, branching points, and network length. We postulate that *ccbe1* has a conserved function in different vascular beds, as previous work showed *ccbe1* to be required for zebrafish lymphangiogenesis and venous differentiation ^34^, as well as murine coronary artery development ^35^.

Our data show that VEGF inhibition decreases vascular volume, surface area, branching points, length and radius, while Notch inhibition increases vascular volume and surface area. This is in agreement with the literature that VEGF is pro-angiogenic, while Notch is angiogenesis limiting ^33,54,55^, and our work provides the first quantitative insights into the extent of this.

The lack of NOS inhibition induced vascular changes was surprising, and future work might examine whether this was due to dose-dependency, whether effects might be observed more locally, or whether side-effects of L-NAME are a contributing factor to this.

The lack of changes in vascular topology upon Wnt inhibition and activation were likely due to the fact that only short-term exposure was performed, and future work might include examining mutants, long-term chemical treatments, and studying dose-dependency.

Our data show that inhibition of F-actin polymerization or Myosin II both result in a reduction of vascular parameters, and we anticipate that this is mainly caused by a mechanical collapse of the cytoskeleton and subsequently the vascular lumen.

The unchanged vascular topology upon glucose treatment is in agreement with previous work, which showed that 24h treatment has no significant effect on vascular topology ^48^.

Additionally, the unchanged vascular topology upon DMSO-treatment, suggests that DMSO indeed only leads to transient vascular changes as previously suggested^50^. These findings clearly demonstrate that quantification of 3D vascular parameters enables meaningful insights into the effects of a range of manipulations on vascular anatomy.

We developed an automated approach to study the vasculature in defined regions, illustrating the approaches utility in the midbrain and hindbrain, as well as for comparison of the left-to-right vasculature. However, the implementation of our analysis approach is not limited to this, allowing the user to select any region of interest.

Future studies are needed to examine vascular symmetry between the left and right vasculature as well as examine stochasticity rules of vascular patterning in more depth, to assess local differences and how these might relate to structural and/or functional brain asymmetry which is established at 4dpf ^56^. We anticipate that our quantification approach will be feasible to allow examination of left-right symmetry in older zebrafish which may be more likely to show vascular asymmetry. Our quantification was performed at the level of the entire cerebral vasculature, and was sufficient to detect significant differences during development and in response to manipulations. Future work might examine vascular parameters in older zebrafish, other transgenic lines, and vessel-specific quantification, which will require an in-depth annotation.

To allow application of the analysis approach by other researchers, we implemented a graphical user interface (GUI; **Fig. S19**) and workflow documentation (**SDoc 1**).

## Conclusion

We here present the first comprehensive workflow to assess zebrafish 3D cerebrovascular similarity, topology, and regional variability. Our data show that robust image analysis is needed to allow objective and quantitative insights. Our results also re-emphasize that the cerebral vascular bed is highly complex and that many mechanisms regulate its formation and patterning.

## Material and Methods

### Zebrafish strains, handling and husbandry

Experiments performed at the University of Sheffield conformed to UK Home Office regulations and were performed under Home Office Project Licence 70/8588 held by TJAC. Maintenance of adult zebrafish in the fish facilities was conducted according to previously described husbandry standard protocols at 28°C with a 14:10 hours (h) light:dark cycle ^57^. Embryos, obtained from controlled pair- or group-mating, were incubated in E3 buffer (5mM NaCl, 0.17mM KCl, 0.33mM CaCl_2_, 0.33mM MgSO_4_) with or without methylene blue. All experiments were conducted with the transgenic reporter line *Tg(kdrl:HRAS-mCherry)^s916^* ^27^.

### Data for biological application

Data were obtained, as described in Kugler et al.^58^. Briefly, the impact of nine chemical components and six morpholinos (Genetools, LLC) on the vascular architecture were examined. Morpholino injections were performed at 1-cell stage using phenol red as injection marker.

Chemical components:

- VEGF inhibition (AV951 250nM 2h ^19^) 4dpf, control n=22, treated n=23, 3 experimental repeats.
- Notch inhibition (DAPT 50µm 12h ^20^) 4dpf, control n=24, treated n=24, 3 experimental repeats.
- Wnt activation (GSK3 inhibitor 10µM 4h ^23^) 3dpf, control n=22, treated n=21, 3 experimental repeats.
- Wnt inhibition (XAV939 10µM 4h ^22^) 3dpf, control n=22, treated n=21, 3 experimental repeats.
- NOS inhibition (LNAME 0.5mM 18h ^21^) 4dpf, control n=16, treated n=17, 2 experimental repeats.
- Myosin II inhibition (Blebbistatin 25µM 1h ^25^) 4dpf, control n=21, treated n=23, 3 experimental repeats.
- F-Actin Polymerization inhibition (Latrunculin B 100nM 1h ^24^) 3dpf, control n=13, treated n=12, 2 experimental repeats.
- Osmotic pressure (Glucose 40nM 24h) 4dpf, control n=18, treated n=18, 2 experimental repeats.
- Membrane Rigidity (DMSO 0.25% 24h) 4dpf, control n=24, treated n=22, 3 experimental repeats.

Morpholinos:

- cardiac troponin T2a (*tnnt2a;* 1.58ng; Genetools, LLC; sequence 5’- CATGTTTGCTCT GATCTGACACGCA-3’) ^15^, 3dpf uninjected control n=17, control MO n= 18, MO n=15, 2 experimental repeats.
- *jagged-1a* (*jag1a*; 0.1ng; 5’-GTCTGTCTGTGTGTCTGTCGCTGTG-3’; Genetools, LLC) ^16^, 3dpf uninjected control n=11, control MO n=12, MO n=10, 2 experimental repeats.
- *jagged-1b* (*jag1b*; 0.8ng; Genetools, LLC; sequence 5’- CTGAACTCCGTCGCAGAATCATGCC - 3’) ^16^, 3dpf uninjected control n=21, control MO n=21, MO n=21, 3 experimental repeats.
- delta-like ligand 4 (*dll4;* 3ng; Genetools, LLC; sequence 5’ - GAGAAAGGTGAGCCAAGCTGCCATG - 3’) ^16^, 3dpf uninjected control n=23, control MO n=23, MO n=23, 3 experimental repeats.
- *notch 1b* (0.25ng; Genetools, LLC; sequence 5’- GTTCCTCCGGTTACCTGGCATACAG - 3’) ^17^, 3dpf uninjected control n=21, control MO n=21, MO n=21, 3 experimental repeats.
- calcium-binding EGF-like domain 1 (*ccbe1)* at 5ng/embryo (Genetools, LLC; sequence 5’-CGGGTAGATCATTTCAGACACTCTG-3’) ^18^, 3dpf uninjected control n=22, control MO n=22, MO n=22, 3 experimental repeats.
- Control MO injection was performed according to the same protocol, with final concentrations as above (5’-CCTCTTACCTCAGTTATTTATA-3’; Genetools, LLC).

### Image Acquisition

Datasets were obtained using a Zeiss Z.1 light sheet microscope with a Plan-Apochromat 20x/1.0 Corr nd=1.38 objective and a scientific complementary metal-oxide semiconductor (sCMOS) detection unit. Data were acquired with activated pivot scan, dual-sided illumination and online fusion; properties of acquired data are as follows: 0.7x zoom, 16bit image depth, 1920 x 1920px (approximately 0.33 x 0.33 µm) image size and minimum z-stack interval (approximately 0.5µm), 561nm laser, LP560, and LP585. Zebrafish embryos were embedded in 2% LMP-agarose containing 200 mg/l Tricaine (MS-222, Sigma). The image acquisition chamber was filled with E3 plus Tricaine (200mg/ml) and maintained at 28°C.

### Image Analysis

Images were analysed using open-source software Fiji ^26^. Data analysis was performed without blinding.

#### Pre-processing and Segmentation

Data were pre-processed performing .czi to .tiff conversion and generation of Maximum Intensity Projections (MIPs) using Bio-Formats Plugin.

Intra-stack motion correction was performed based on scale invariant feature transform SIFT ^59,60^ using the Fiji Plugin “Linear Stack Alignment with SIFT” with parameters as described previously ^10,11^.

Vascular enhancement was performed assuming local vessel tubularity using the Fiji Sato enhancement filter “Analyse>Tubeness” ^28^ with an optimised parameter set as described previously ^10,11,14^. Enhanced images were segmented using Otsu thresholding ^29^ as described in ^10,11^.

#### Manual measurements

The following measurements were performed manually using Fiji line region of interest (ROI) tool and “Analyse>Measure”. Growth expansion of the primordial midbrain channel (PMBC) was measured at the inner vascular edge at the position behind eyes. Distance of middle metencephalic central artery (MMCtA) was measured at the most anterior point. MMCtA angle was measured in the right MMCtA from its posterior to its anterior end. Posterior metencephalic central artery (PMCtA) angle was measured from its origin in the centre of the embryo to its right lateral end.

#### Inter-sample Registration

Visual comparison of samples was used to identify and extract 11 anatomical landmarks which were found in all samples and distributed along the anterior-posterior, left-right, and dorso-ventral body axis. These 11 anatomical landmarks (**Fig. 2**) were then used for manual landmark-based rigid registration using the Fiji Plugin “Name Landmarks and Register”. The automatic rigid inter-sample registration method, by Johannes Schindelin and Mark Longair, based on the Virtual Insect Brain Protocol (VIB) ^30,31^, was used with the following parameters: no initial transformation, 5 best matching orientations for further optimization, image down-sampling 2-6 times, no ROI reduction (*i.e.* bounding box selection) for optimisation, using Euclidean measure of difference and showing transformed image.

Target embryos, used as a template for registration, were selected based on **(a)** sample body-axis orientation along image axis (anterior-posterior along image y-axis, coronal plane along image z-axis and x-axis), **(b)** all major vessels visualized in image, **(c)** no obvious abnormalities (*ie.* vascular patterning visually normal, no haemorrhaging, no gross defects).

A dataset for registration validation was constructed by repetitive image acquisition of the same sample after manual displacement. This allowed quantification of registration outcomes following registration to the initial acquisition.

To further examine 3D topological similarity between samples, six embryos were registered to a target and averages extracted.

#### Similarity assessment

Vascular intra-sample left-right and inter-sample similarity (S_n_) were visually assessed and Dice coefficient between target (T) and moving image (M) quantified using the Fiji Plugin MorphoLibJ ^61^ (Eq.1).

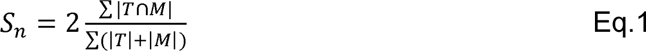

#### Quantification

Vascular volume (V_n_; Eq.2) quantification was performed in a cranial ROI as described previously ^10,11^, by multiplying histogram black voxel count (V_black_) following segmentation by the respective voxel volume (V_x,y,z_).

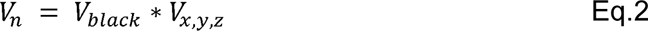

Vascular surface voxels (S_n_) were quantified in the same cranial ROI after edge detection using Canny Edge detection ^62^.

Vascular density (D_n_; Eq.3) was quantified as vascular volume (V_n_) as a proportion of total cranial ROI volume (Vtotal).

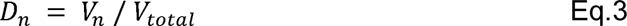

Next, data were down-sampled (1920px to 512px using bilinear interpolation) to decrease processing time in the following steps:

Euclidean Distance Maps (EDM_n_) of vascular voxel (v) distance to the nearest background voxel (b) were produced from binary segmented images using the Fiji plugin “Distance Map 3D”, which calculates distance in three-dimensional Euclidean space (Eq.4; Process > Binary > Distance Map in 3D; implemented by Jens Bache- Wiig and Christian Henden^63^).

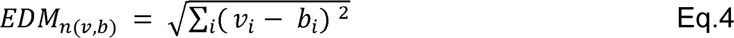

Centreline extraction was performed in segmented images using the Fiji “Skeletonize 2D/3D” Plugin (by Ignacio Arganda-Carreras), based on 3D thinning ^64^, using a layer-by-layer removal. Centreline voxels (zero-valued in images) were quantified for total network length (L_n_) analysis by quantification of vascular voxels (zero value) in histogram.

The “Analyse Skeleton” Plugin in Fiji (Analyse > Skeleton > Analyse Skeleton 2D/3D; implemented by Ignacio Arganda-Carreras ^65^) was used to identify and measure the number of branching points (Analyse > Skeleton > Summarize Skeleton).

To quantify vessel radii (R_n_), EDMs were multiplied with extracted skeletons, resulting in 1D representation of vessel radii as represented by intensity of voxels.

To analyse vascular complexity (C_n_) by assessment of branching, Sholl dentritic arbor analysis (Eq.5) ^66,67^ was applied to 2D skeleton MIPs using the Fiji Sholl Analysis plugin ^68^ (developed and maintained by Tom Maddock, Mark Hiner, Curtis Rueden, Johannes Schindelin, and Tiago Ferreira).

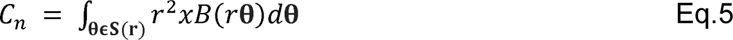

Where C_n_ is vascular complexity given by Sholl intersection profiles as number of branches with the distance from the shell centre, calculated with spheres of radius S(r), B being the region of interest examined, θ being a point on S(r), and χB(r,θ) is the function for point (r,θ in B. Default parameters were used to create a sphere, except for distance from centre (700 voxel being the sphere size) and step size (5 voxel being the shell step-size). The centre of the sphere was placed in the midbrain at the BA-PCS junctions.

#### Intra-sample Symmetry

Intra-sample symmetry was assessed using the segmented vasculature to allow topological rather than intensity-based assessments. After manual selection of the central body axis (centrally along the basal aorta (BA), between L-R posterior communicating segments (PCS), between L-R MMCtA and the bifurcation point of L-R anterior cerebral vein (ACeV)), automatic image rotation was performed to align the anterior-posterior axis of samples with the image y-axis to allow for mirroring. Following, the right vascular volume was automatically mirrored by horizontal image transformation and similarity was derived via voxel-overlap.

Data representation

To visualize data, maximum intensity projections (MIP) were generated and intensity inversion was applied as appropriate to give the clearest rendering of relevant structures.

### Statistical analysis

Normality of data was tested using D’Agostino-Pearson omnibus test. Statistical analysis of normally distributed data was performed using a One-way ANOVA to compare multiple groups or Student’s t-test to compare two groups. Non-normally distributed data were analysed with a Kruskal-Wallis test to compare multiple groups, or Mann-Whitney test to compare two groups. Analysis was performed in GraphPad Prism Version 7 (GraphPad Software, La Jolla California USA). P values are indicated as follows: p<0.05 *, p<0.01 **, p<0.001 ***, p<0.0001 ****. Data represents mean and standard deviation (s.d.), if not otherwise stated. Cluster analysis was performed using ClustVis ^69^. Image representation was performed using Inkscape (https://www.inkscape.org). Data are available on request.

## Author contributions

Funding obtained by PAA and TJAC; Data Acquisition, ECK, VS, KP, KC; Investigation, Validation, and Data Curation, ECK and JLTF; Formal Visualization and Analysis, ECK; Resources, PA and TC; Project Administration, ECK; Writing – Original Draft, ECK; Writing – Review and Editing, all authors.

## Acknowledgement

We thank the funders who supported this work, including the Department of Infection, Immunity and Cardiovascular Disease (University of Sheffield) and the Insigneo Institute for in silico medicine (University of Sheffield) awarding funding to ECK, and the British Heart Foundation for Infrastructure Award IG/15/1/31328 awarded to TJAC. We are grateful to Steven Renshaw and Freek van Eeden for their helpful feedback on the manuscript. We thank Robert N. Wilkinson, Stefan Schulte- Merker, Emily Noël, the Perak lab Sheffield, and Henry Roehl for sharing chemical compounds. The authors thank the Bateson Centre aquarium facility for zebrafish husbandry.

**SDoc 1: Workflow Documentation for using the image analysis approach (.pdf file).**

Deposited on Github (https://github.com/ElisabethKugler/ZFVascularQuantification) and doi assigned with zenodo (https://doi.org/10.5281/zenodo.3978278).

**SDoc 2: Code – Individual Macros and code for GUI (.zip file).**

Deposited on Github (https://github.com/ElisabethKugler/ZFVascularQuantification) and doi assigned with zenodo (https://doi.org/10.5281/zenodo.3978278).

**SDoc 3: Test Data.**

**SDoc 4: Screencast.**

## Supplementary Figures

**Figure S1:**
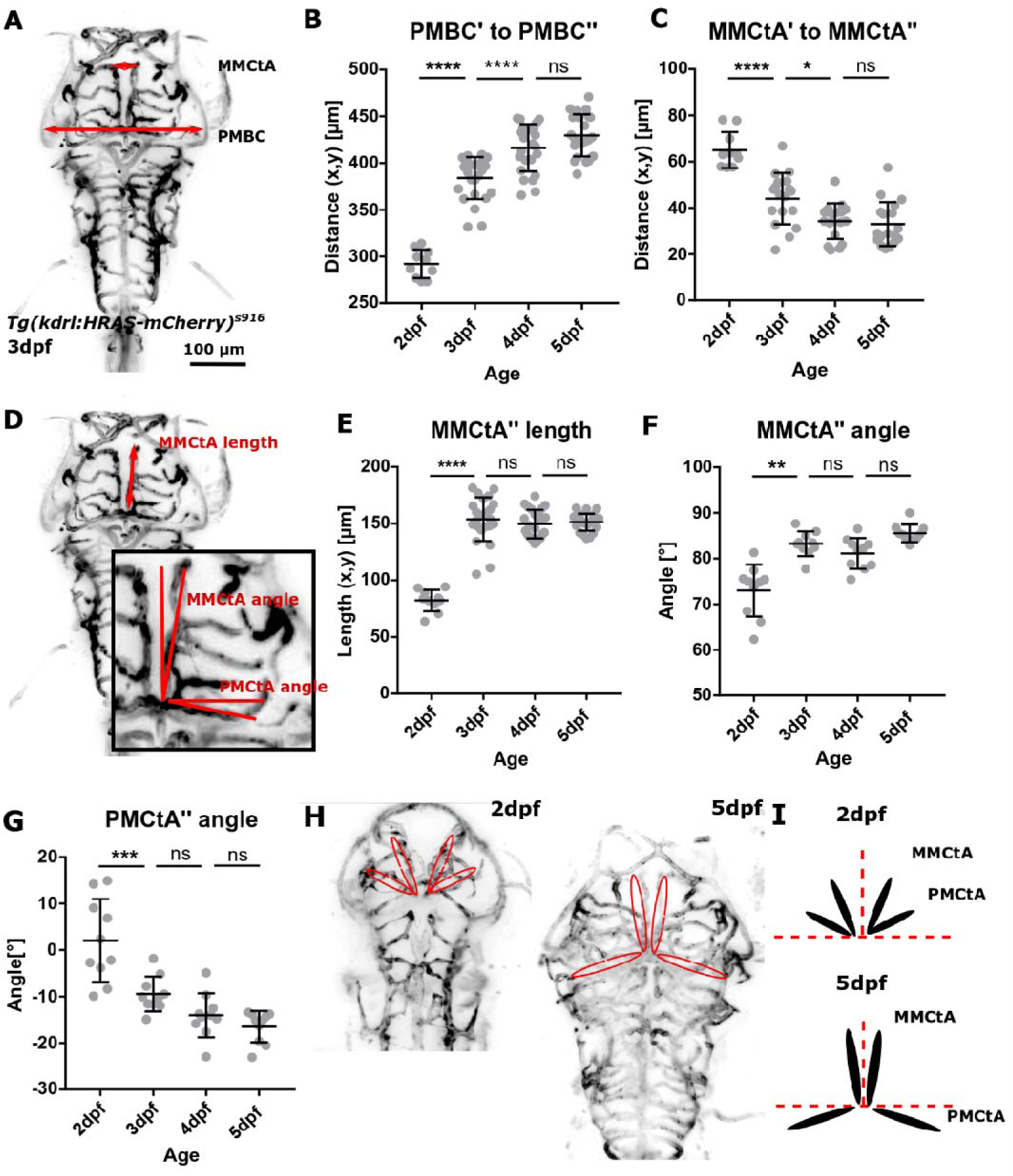
Manual quantification allowed the identification of anatomical landmarks for registration. **A** Manual measurements were performed on the primordial midbrain channel (PMBC) and middle mesencephalic central artery (MMCtA). **B** Distance between PMBC’ and PMBC’’ was found to statistically significantly increase from 2-to-4dpf (2-3dpf p<0.0001, 3-4dpf p<0.0001, 4-5dpf p 0.1786; embryos n 2dpf=10 (2 experimental repeats), 3dpf=25, 4dpf=25, 5dpf=25; 4 experimental repeats; One-way ANOVA). **C** Distance between anterior ends of MMCtA’ and MMCtA’’ was found to statistically significantly decrease from 2-to-4dpf (2-3dpf p<0.0001, 3-4dpf p 0.0137, 4-5dpf p 0.9739; embryos n 2dpf=10 (2 experimental repeats), 3dpf=18, 4dpf=18, 5dpf=18; 3 experimental repeats; One-way ANOVA). **D** Manual measurements were performed to measure MMCtA’’ length, MMCtA’’angle, and PMCtA’’ angle. **E** MMCtA’’ length was found to statistically significantly increase from 2-to-3dpf, but not 3-to-5dpf (2-3dpf p<0.0001, 3-4dpf p 0.7138, 4-5dpf p 0.9773; embryos n 2dpf=10 (2 experimental repeats), 3dpf=24, 4dpf=24, 5dpf=24; 4 experimental repeats; One-way ANOVA). **F** MMCtA’’ angle was found to statistically significantly increase from 2-to-3dpf, but not 3-to-5dpf (2-3dpf p 0.0028, 3-4dpf p>0.9999, 4-5dpf p 0.0657; embryos n 2dpf=10, 3dpf=10, 4dpf=10, 5dpf=10; 2 experimental repeats; Kruskal-Wallis test). **G** PMCtA’’ angle was found to statistically significantly decrease from 2-to-3dpf, but not 3-to-5dpf (2-3dpf p 0.0003, 3-4dpf p 0.2790, 4-5dpf p 0.7652; embryos n 2dpf=10, 3dpf=10, 4dpf=10, 5dpf=10; 2 experimental repeats; One-way ANOVA). **G,H** Data showed that MMCtA moved towards embryonic midline, whilst PMCtA moved posteriorly.

**Figure S2:**
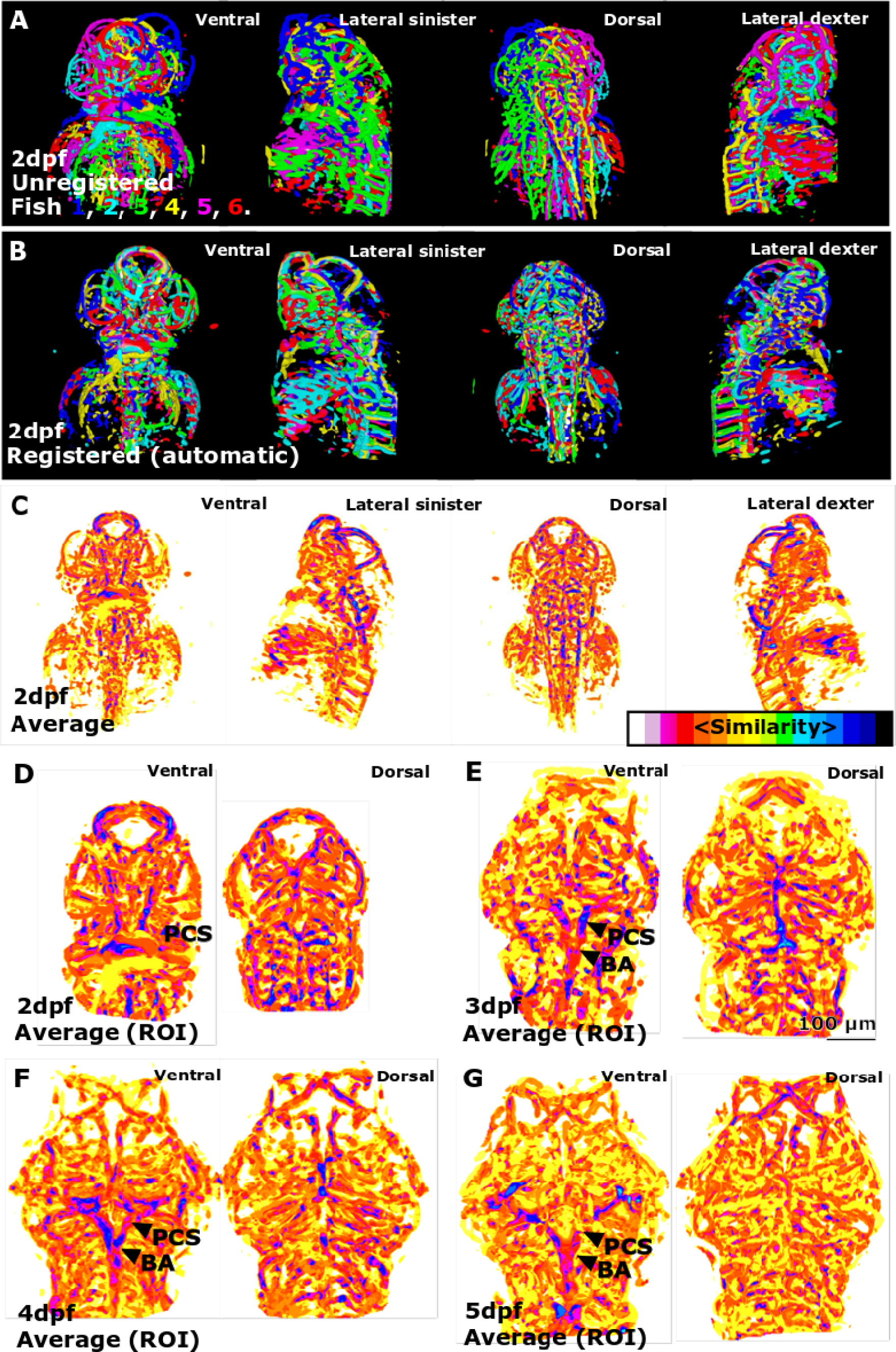
Sample registration allows visualisation of inter-individual vascular similarity. **(A)** 3D rendering of six unregistered segmented embryos with individual embryos
being colour-coded. **(B)** 3D rendering of the same six embryos as (A) following registration. **(C)** 3D rendering of the same six embryos as (B) following averaging, showing regions of increased similarity in darker colours. **(D)** 3D rendering of the same six embryos as (C) following ROI selection. **(E)** 3D rendering of six 3dpf embryos following segmentation, registration, averaging, and ROI selection. **(F)** 3D rendering of six 4dpf embryos following segmentation, registration, averaging, and ROI selection. **(G)** 3D rendering of six 5dpf embryos following segmentation, registration, averaging, and ROI selection.

**Table S1:**
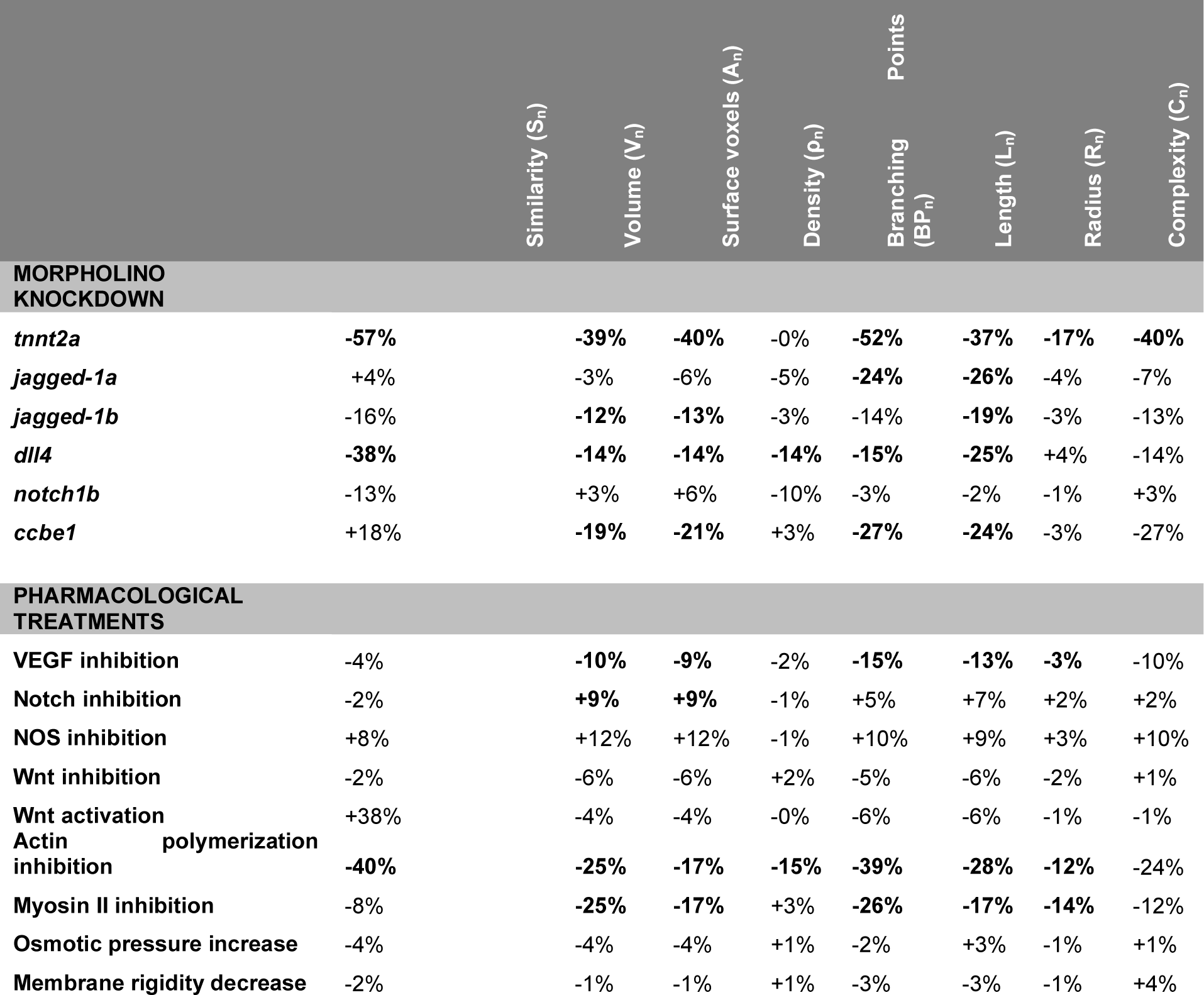
Quantification of similarity, volume, surface voxels, density, branching points, network length, network radius, and network complexity was conducted following morpholino-based knock-down of *tnnt2a, jagged-1a, jagged-1b, dll4, notch1b*, and *ccbe1* and application of chemicals for VEGF inhibition, Notch inhibition, NOS inhibition, Wnt inhibition, Wnt activation, F-actin polymerization inhibition, Myosin II inhibition, osmotic pressure (OP) increase, and membrane rigidity (MR) decrease. Percentages represent difference of mean value between control morpholino (MO) and MO, or controls and treatment groups. Bold indicates statistically significant difference. *Abbreviations: act. – activation, inh. – inhibition, NOS – nitric oxide synthase, MR – membrane rigidity, OP – osmotic pressure;*

**Figure S3:**
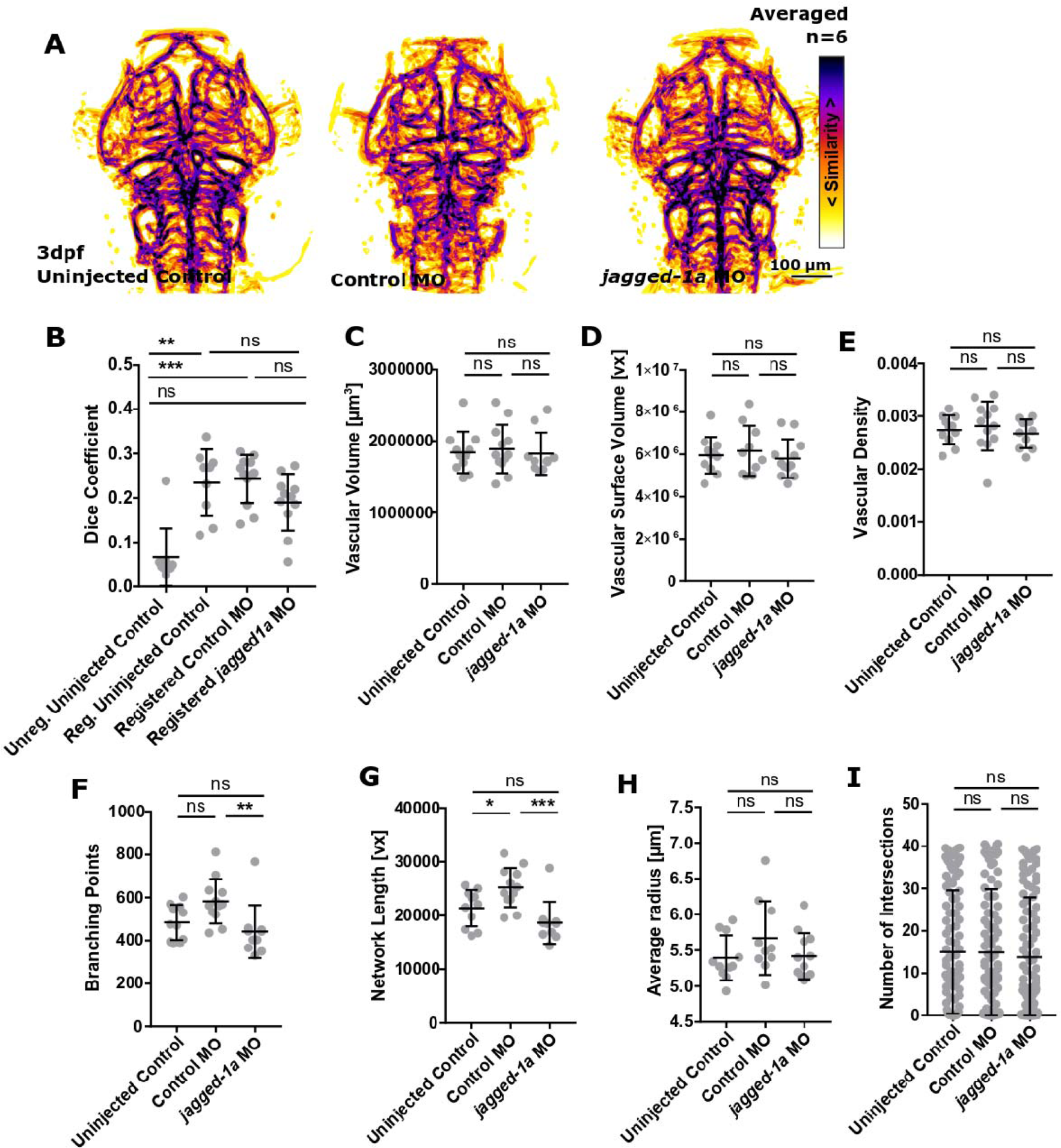
The impact *jagged-1a* knock-down on vascular topology. **A** MIPs of averaged data of uninjected controls, control MO, and *jagged-1a* MO following segmentation and registration. **B** No statistically significant difference was found when comparing registered control MO to *jagged-1a* MO (p 0.6359; uninjected control=9, control MO=12, *jagged-1a* MO=10; Kruskal-Wallis test). **C** Vascular volume was not statistically significantly changed in *jagged-1a* MO (uninjected control p>0.9999, control MO p>0.9999; uninjected control=11, control MO=12, *jagged-1a* MO=10; Kruskal Wallis test). **D** Vascular surface was not statistically significantly changed in *jagged-1a* MO (uninjected control p 0.9254, control MO p 0.6702; One-Way ANOVA). **E** Vascular density was not statistically significantly changed in *jagged-1a* MO (uninjected control p 0.8905, control MO p 0.6131; One-Way ANOVA). **F** Branching points were statistically significantly decreased in *jagged-1a* MO in comparison to control MO (uninjected control p 0.6219, control MO p 0.0041; Kruskal-Wallis test). **G** Vascular network length was statistically significantly decreased in *jagged-1a* MO in comparison to control MO (uninjected control p 0.2140, control MO p 0.0006; One-Way ANOVA). **H** Average vessel radius was not statistically significantly changed in *jagged-1a* MO (uninjected control p 0.9876, control MO p 0.2966; One-Way ANOVA). **I** Vascular complexity was not statistically significantly changed in *jagged-1a* MO (uninjected control p>0.9999, control MO p>0.9999; Kruskal-Wallis test).

**Figure S4:**
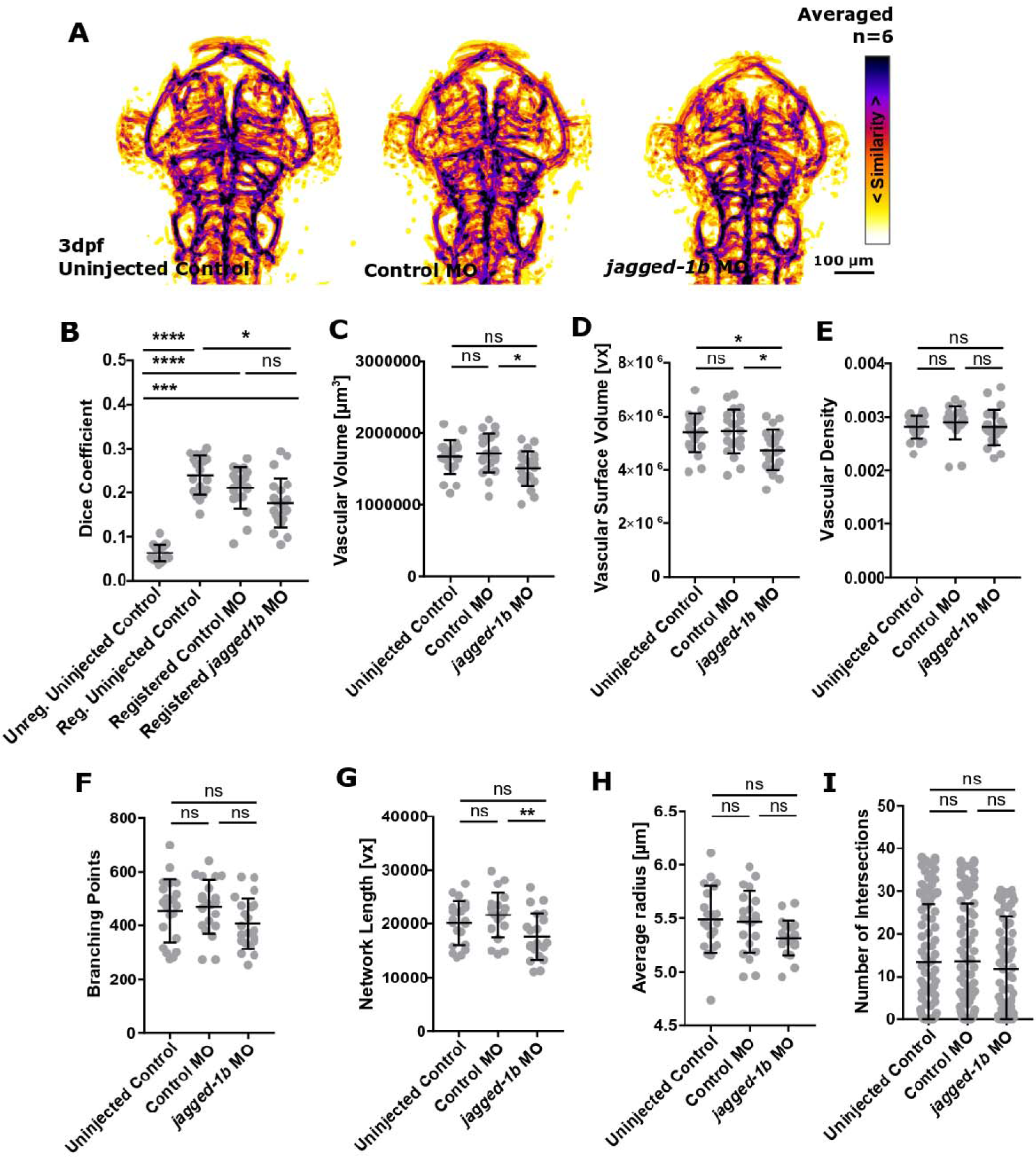
The impact *jagged-1b* knock-down on vascular topology. **A** MIPs of averaged data of uninjected controls, control MO, and *jagged-1b* MO following segmentation and registration. **B** No statistically significant difference was found when comparing registered control MO to *jagged-1b* MO (p 0.4737; uninjected control=18, control MO=21, *jagged-1b* MO=21; Kruskal-Wallis test). **C** Vascular volume was statistically significantly decreased in *jagged-1b* MO in comparison to control MO (uninjected control p 0.0848, control MO p 0.0222; uninjected control=21, control MO=21, *jagged-1b* MO=21; One-Way ANOVA). **D** Vascular surface was statistically significantly decreased in *jagged-1b* MO (uninjected control p 0.0204, control MO p 0.0124; One-Way ANOVA). **E** Vascular density was not statistically significantly changed in *jagged-1b* MO (uninjected control p>0.9999, control MO p 0.1694; One-Way ANOVA). **F** Branching points were not statistically significantly changed in *jagged-1b* MO (uninjected control p 0.2515, control MO p 0.1484; One-Way ANOVA). **G** Vascular network length was statistically significantly decreased in *jagged-1b* MO in comparison to control MO (uninjected control p 0.1339, control MO p 0.0082; One-Way ANOVA). **H** Average vessel radius was not statistically significantly changed in *jagged-1b* MO (uninjected control p 0.0922, control MO p 0.1601; One-Way ANOVA). **I** Vascular complexity was not statistically significantly changed in *jagged-1b* MO (uninjected control p >0.9999, control MO p 0.7584; Kruskal-Wallis test).

**Figure S5:**
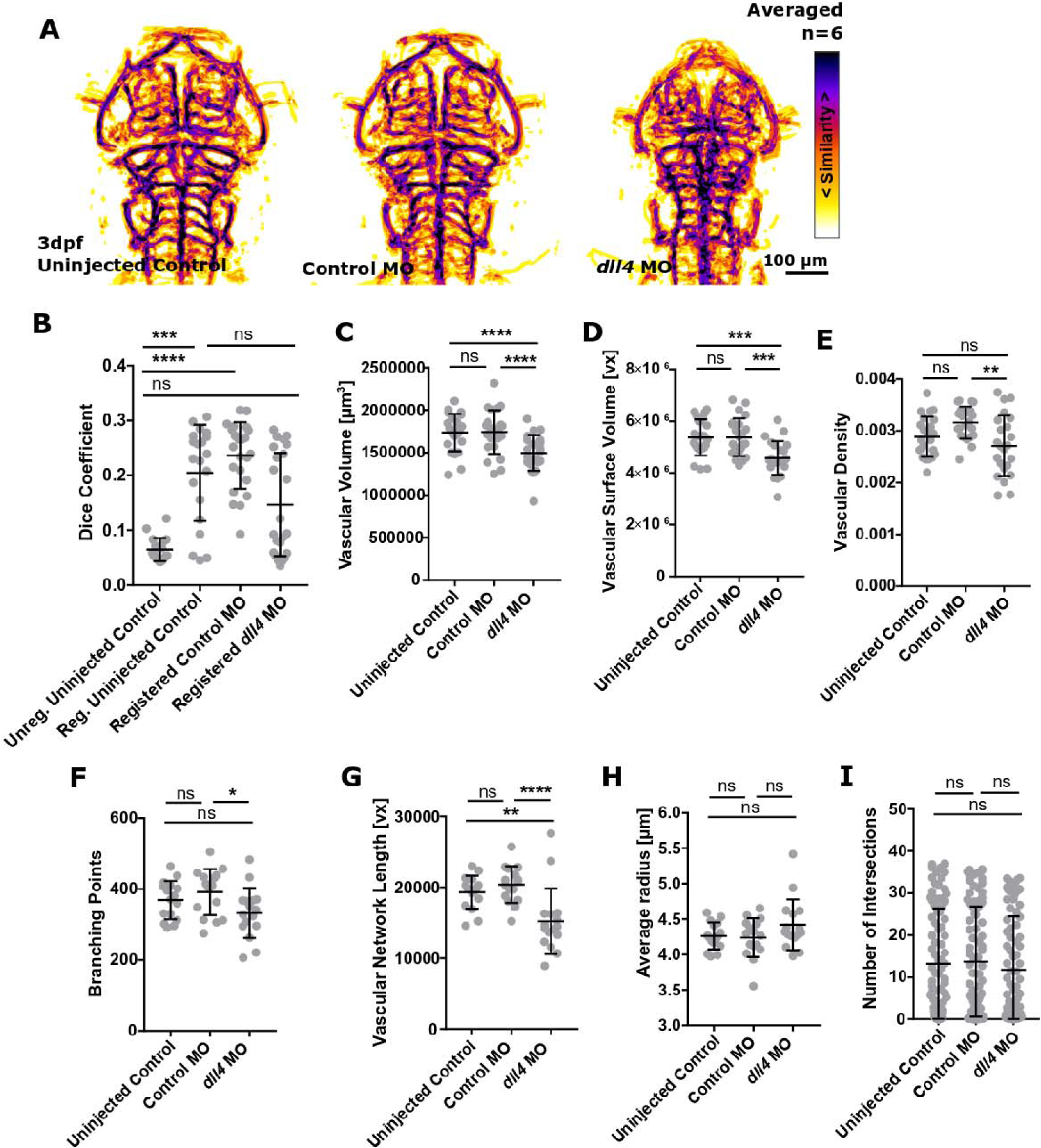
The impact *dll4* knock-down on vascular topology. **A** MIPs of averaged data of uninjected controls, control MO, and *dll4* MO following segmentation and registration. **B** A statistically significant decrease was found when comparing registered control MO to *dll4* MO (p 0.0103; uninjected control=20, control MO=23, *dll4* MO=23; Kruskal-Wallis test). **C** Vascular volume was statistically significantly decreased in *dll4* MO (uninjected control p 0.0023; control MO p 0.0017; uninjected control=23, control MO=23, *dll4* MO=23; One-Way ANOVA). **D** Vascular surface was statistically significantly decreased in *dll4* MO (uninjected control p 0.0007, control MO p 0.0006; One-Way ANOVA). **E** Vascular density was statistically significantly decreased in *dll4* MO in comparison to control MO (uninjected control p 0.3718, control MO p 0.0028; One-Way ANOVA). **F** Branching points were statistically significantly decreased in *dll4* MO in comparison to control MO (uninjected control p 0.4662, control MO p 0.0484; Kruskal-Wallis test). **G** Vascular network length was statistically significantly decreased in *dll4* MO (uninjected control p 0.0076, control MO p 0.0004; Kruskal-Wallis test). **H** Average vessel radius was not statistically significantly changed in *dll4* MO (uninjected control p 0.5547, control MO p 0.7673; Kruskal-Wallis test). **I** Vascular complexity was not statistically significantly changed in *dll4* MO (uninjected control p>0.9999, control MO p>0.9999; Kruskal-Wallis test).

**Figure S6:**
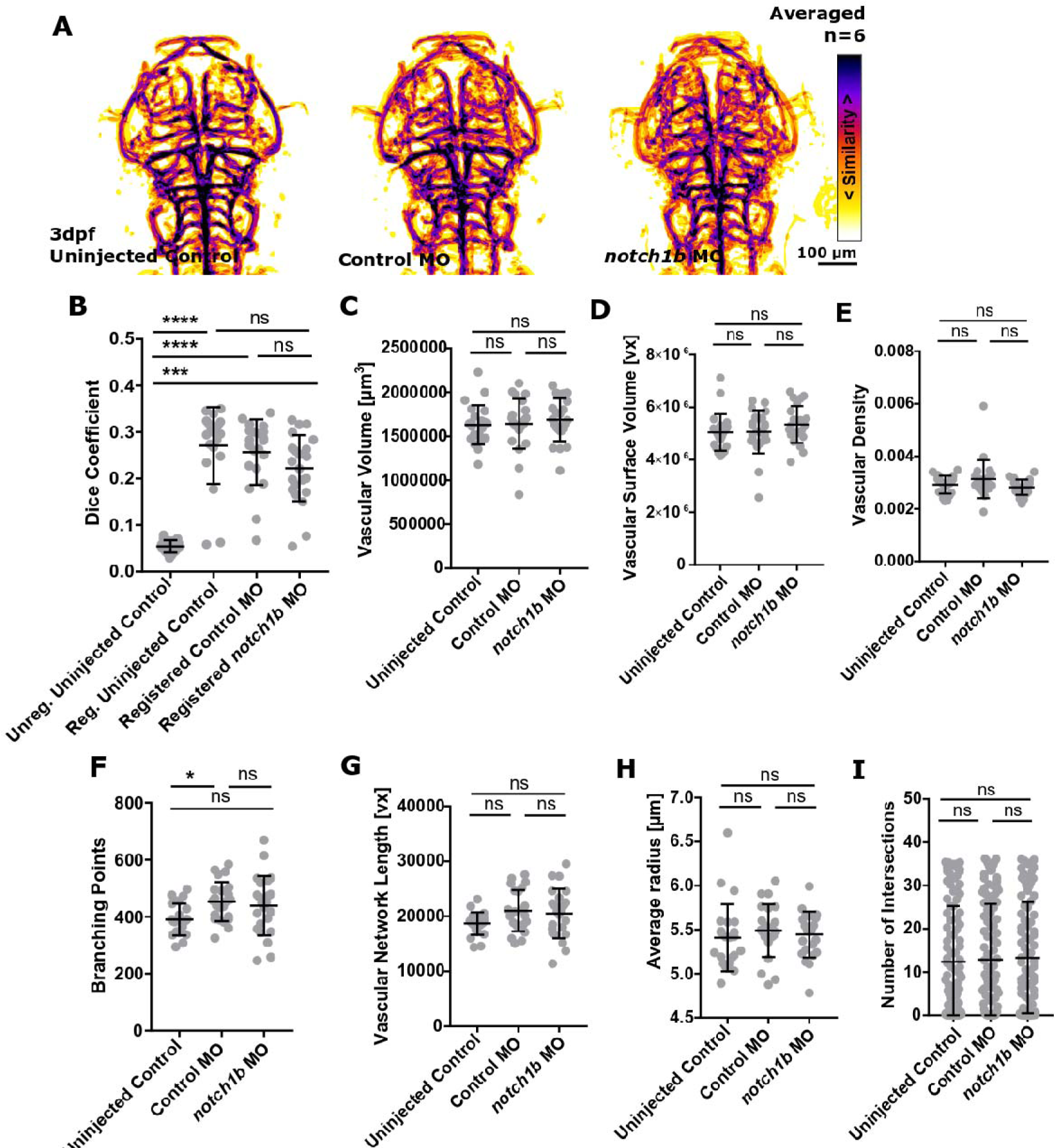
The impact *notch1b* knock-down on vascular topology. **A** MIPs of averaged data of uninjected controls, control MO, and *notch1b* MO following segmentation and registration. **B** No statistically significant difference was found when comparing registered control MO to *notch1b* MO (p 0.9922; uninjected control=20, control MO=21, *notch1b* MO=23; Kruskal-Wallis test). **C** Vascular volume was not statistically significantly changed in *notch1b* MO (uninjected control p 0.9634, control MO p>0.9999; uninjected control=23, control MO=23, *notch1b* MO=23; Kruskal-Wallis test). **D** Vascular surface was not statistically significantly changed in *notch1b* MO (uninjected control p 0.2296, control MO p>0.9999; Kruskal-Wallis test). **E** Vascular density was not statistically significantly changed in *notch1b* MO (uninjected control p 0.6589, control MO p 0.1010; Kruskal-Wallis test). **F** Branching points were not statistically significantly changed in *notch1b* MO (uninjected control p 0.1018, control MO p 0.8225; One-Way ANOVA). **G** Vascular network length was not statistically significantly changed in *notch1b* MO (uninjected control p 0.2077, control MO p 0.8787; One-Way ANOVA). **H** Average vessel radius was not statistically significantly changed in *notch1b* MO (uninjected control p 0.7102, control MO p>0.9999; Kruskal-Wallis test). **I** Vascular complexity was not statistically significantly changed in *notch1b* MO (uninjected control p>0.9999, control MO p>0.9999; Kruskal-Wallis test).

**Figure S7:**
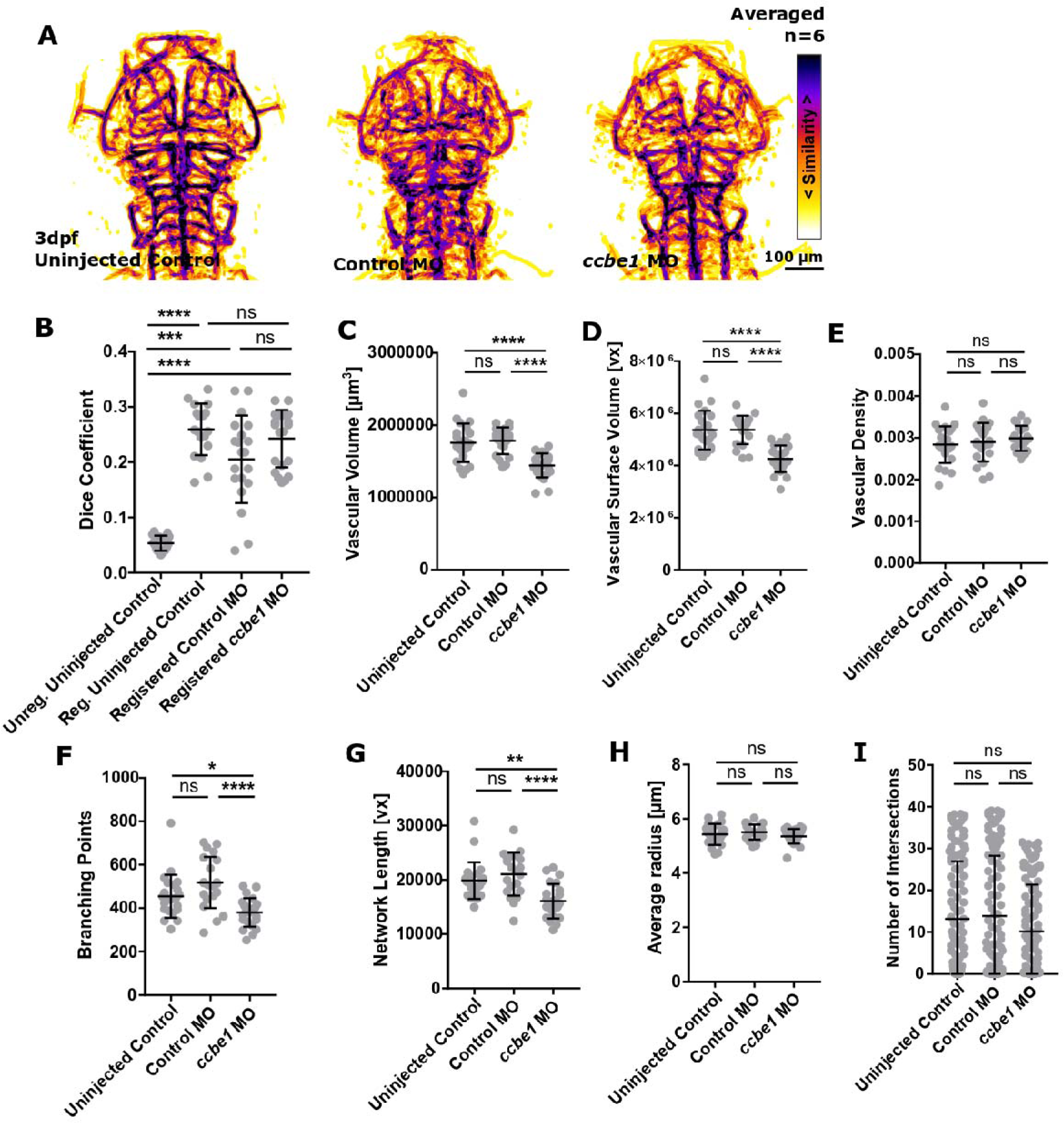
The impact *ccbe1* knock-down on vascular topology. **A** MIPs of averaged data of uninjected controls, control MO, and *ccbe1* MO following segmentation and registration. **B** No statistically significant difference was found when comparing registered control MO to *ccbe1* MO (p 0.9061; uninjected control=19, control MO=20, *ccbe1* MO=21; Kruskal-Wallis test). **C** Vascular volume was statistically significantly decreased in *ccbe1* MO (uninjected control p<0.0001, control MO p<0.0001; uninjected control=23, control MO=21, *ccbe1* MO=23; One-Way ANOVA). **D** Vascular surface was statistically significantly decreased in *ccbe1* MO (uninjected control p<0.0001, control MO p<0.0001; One-Way ANOVA). **E** Vascular density was not statistically significantly changed in *ccbe1* MO (uninjected control p 0.4194, control MO p 0.7287; One-Way ANOVA). **F** Branching points were statistically significantly decreased in *ccbe1* MO (uninjected control p 0.0283, control MO p<0.0001; Kruskal-Wallis test). **G** Vascular network length was statistically significantly decreased in *ccbe1* MO (uninjected control p 0.0032, control MO p<0.0001; Kruskal-Wallis test). **H** Average vessel radius was not statistically significantly changed in *ccbe1* MO (uninjected control p 0.6170, control MO p 0.3152; One-Way ANOVA). **I** Vascular complexity was not statistically significantly changed in *ccbe1* MO (uninjected control p 0.8835, control MO p 0.6544; Kruskal-Wallis test).

**Figure S8:**
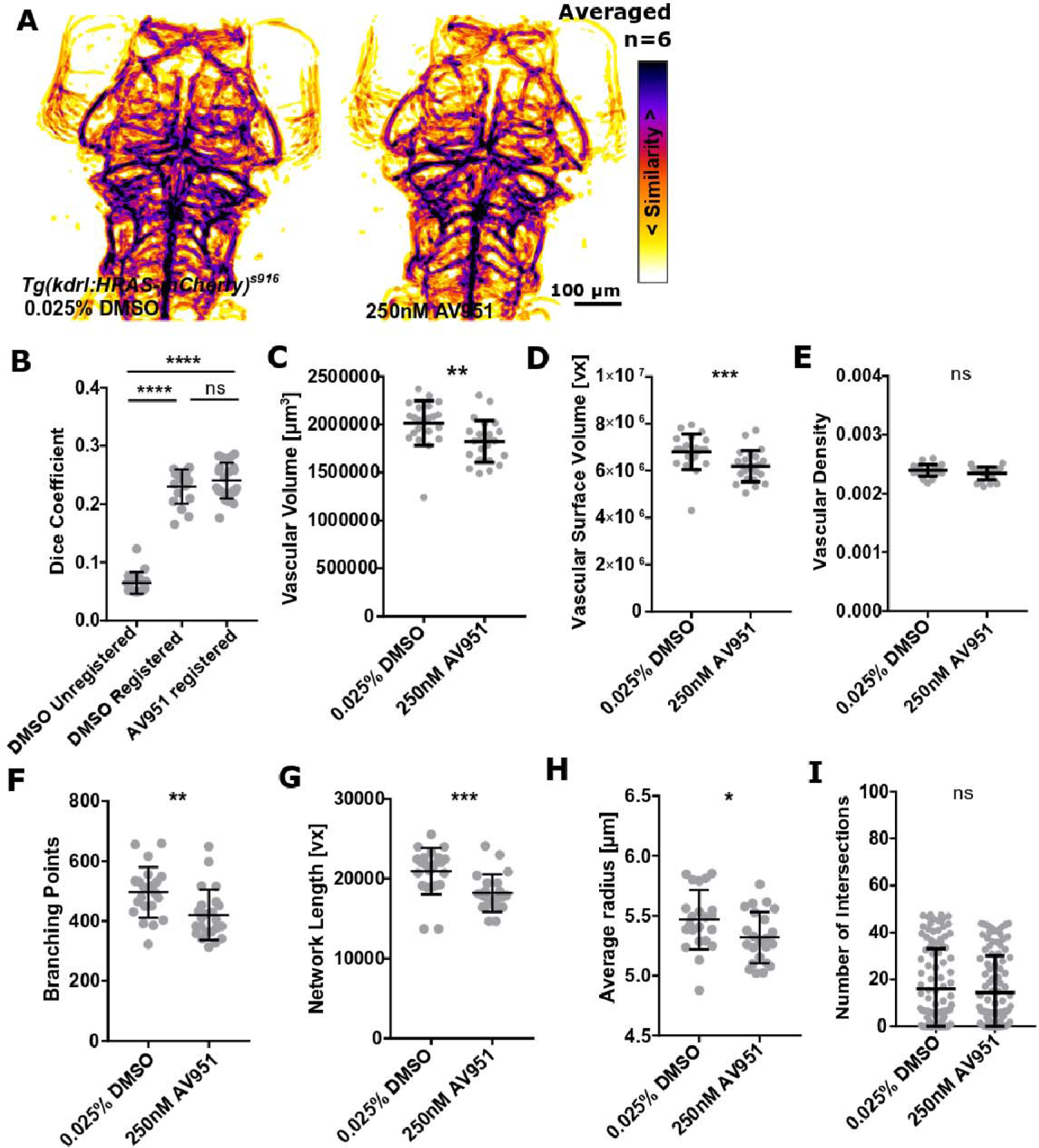
The impact of VEGF inhibition on vascular topology. **A** MIPs of averaged data of control and AV951 treated samples following segmentation and registration. **B** No statistically significant difference was found when comparing registered controls to VEGF inhibitor treated samples (p>0.9999; control=18, AV951=23; Kruskal-Wallis test). **C** Vascular volume was statistically significantly decreased in AV951 treated samples (p 0.0014; control=22, AV951=23; Mann-Whitney U test). **D** Vascular surface was statistically significantly decreased in AV951 treated samples (p 0.0010; Mann-Whitney U test). **E** Vascular density was not statistically significantly changed in AV951 treated samples (p 0.1048; unpaired t-test). **F** Branching points were statistically significantly decreased in AV951 treated samples (p 0.0016; Mann-Whitney U test). **G** Vascular network length was not statistically significantly changed in AV951 treated samples (p 0.0004; Mann-Whitney U test). **H** Average vessel radius was statistically significantly reduced in AV951 treated samples (p 0.0371; unpaired Student’s t-test). **I** Vascular complexity was not statistically significantly changed in AV951 treated samples (p 0.4949; Mann-Whitney U test).

**Figure S9:**
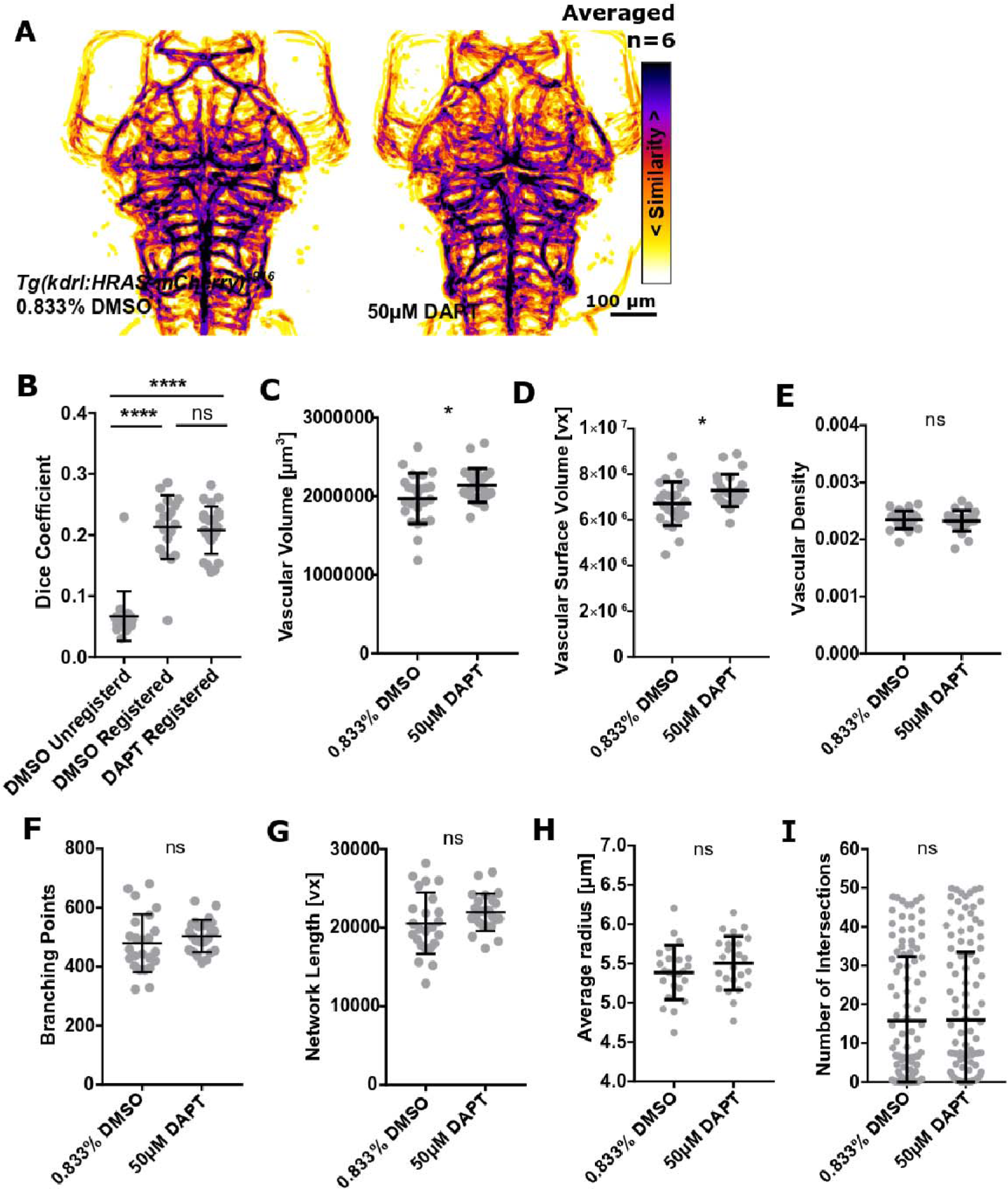
The impact of Notch inhibition on vascular topology. **A** MIPs of averaged data of control and DAPT treated samples following segmentation and registration. **B** No statistically significant difference was found when comparing registered controls to Notch inhibitor treated samples (p>0.9999; control=19, DAPT=24; Kruskal-Wallis test). **C** Vascular volume was statistically significantly increased in DAPT treated samples (p 0.039; control=24, DAPT=24; unpaired t-test). **D** Vascular surface was statistically significantly decreased in DAPT treated samples(p 0.0226; unpaired t-test). **E** Vascular density was not statistically significantly changed in DAPT treated samples (p 0.8023; Mann-Whitney U test). **F** Branching points were not statistically significantly changed in DAPT treated samples (p 0.2976; unpaired Student’s t-test). **G** Vascular network length was not statistically significantly changed in DAPT treated samples (p 0.1436; unpaired Student’s t-test). **H** Average vessel radius was not statistically significantly changed in DAPT treated samples (p 0.2334; unpaired Student’s t-test). **I** Vascular complexity was not statistically significantly changed in DAPT treated samples (p 0.8276; Mann-Whitney U test).

**Figure S10:**
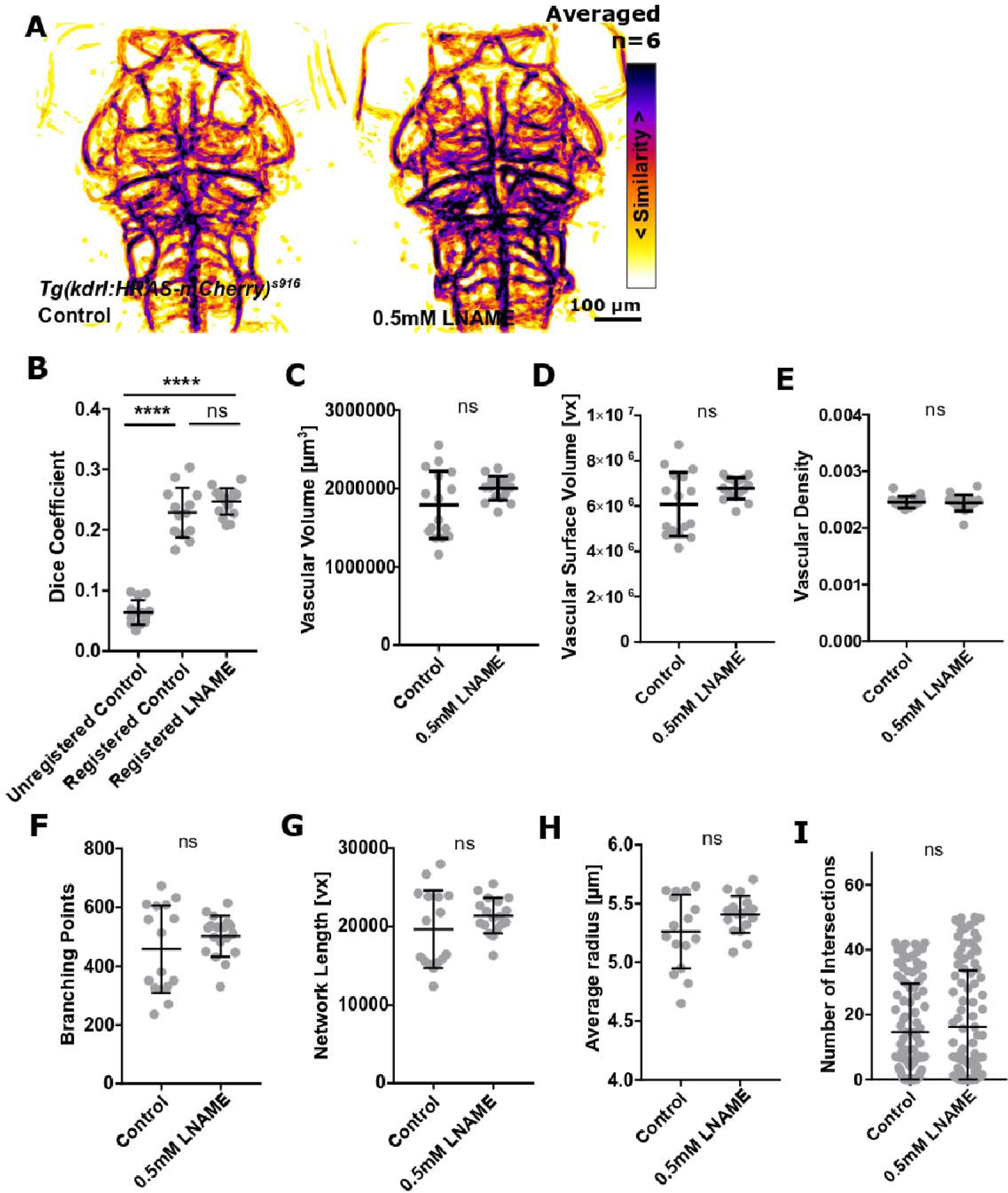
The impact of NOS inhibition on vascular topology. **A** MIPs of averaged data of control and L-NAME treated samples following segmentation and registration. **B** No statistically significant difference was found when comparing registered controls to NOS inhibitor treated samples (p 0.2008; control=13, L-NAME=17; One-way ANOVA). **C** Vascular volume was not statistically significantly changed in L-NAME treated samples (p 0.0642; control=16, L-NAME=17; unpaired t-test). **D** Vascular surface was not statistically significantly changed in L-NAME treated samples (p 0.0600; unpaired t-test). **E** Vascular density was not statistically significantly changed in L-NAME treated samples (p 0.8939; Mann-Whitney U test). **F** Branching points were not statistically significantly changed in L-NAME treated samples (p 0.6272; Mann-Whitney U test). **G** Vascular network length was not statistically significantly changed in L-NAME treated samples (p 0.6272; Mann-Whitney U test). **H** Average vessel radius was not statistically significantly changed in L-NAME treated samples (p 0.0913; unpaired Student’s t-test). **I** Vascular complexity was not statistically significantly changed in L-NAME treated samples (p 0.7850; Mann-Whitney U test).

**Figure S11:**
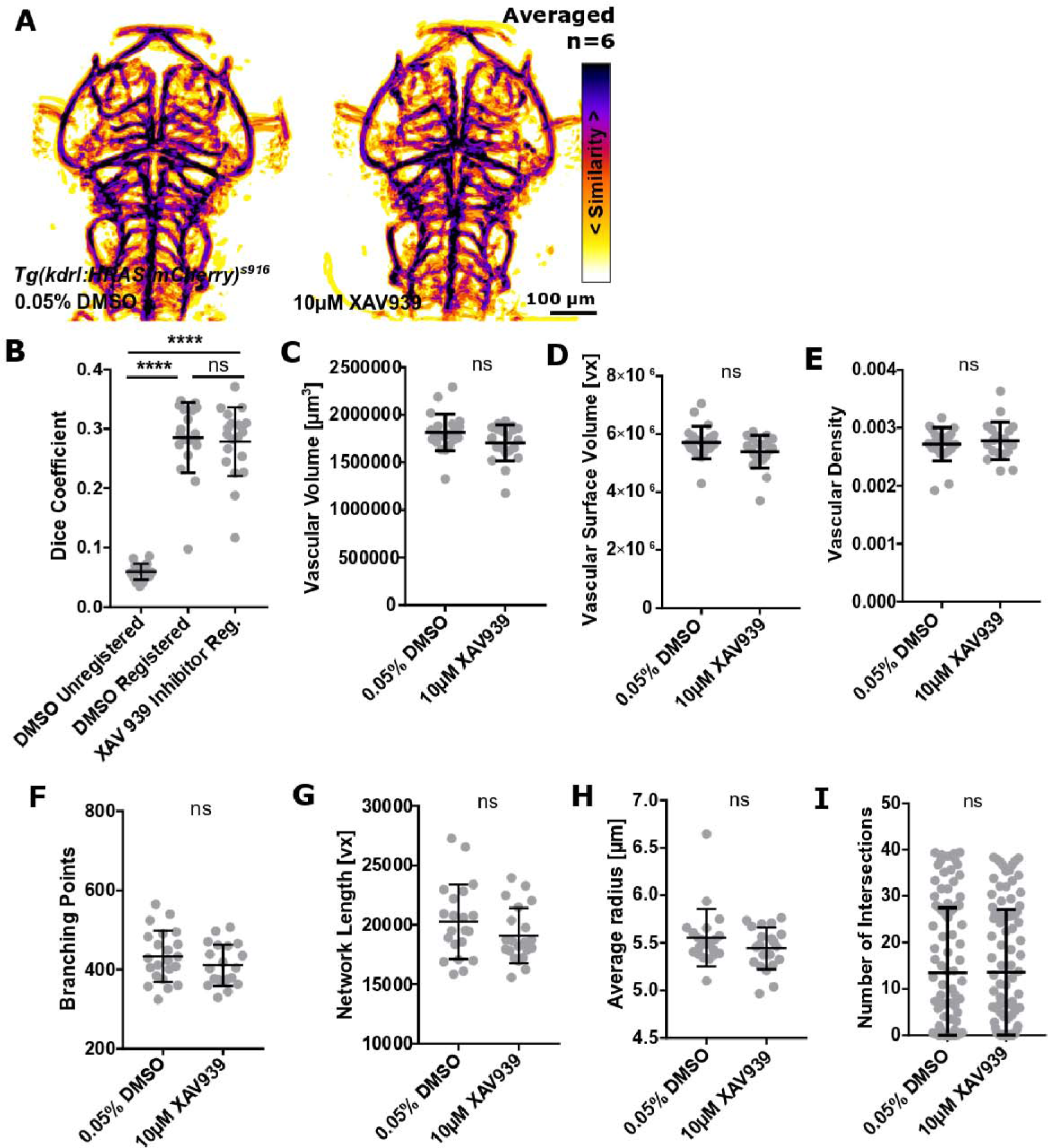
The impact of Wnt inhibition on vascular topology. **A** MIPs of averaged data of control and XAV939 treated samples following segmentation and registration. **B** No statistically significant difference was found when comparing registered controls to Wnt inhibitor treated samples (p>0.9999; control=19, XAV939=20; Kruskal-Wallis test). **C** Vascular volume was not statistically significantly changed in XAV939 treated samples (p 0.1419; control=22, XAV939=20; Mann-Whitney U test). **D** Vascular surface was not statistically significantly changed in XAV939 treated samples (p 0.0994; Mann-Whitney U test). **E** Vascular density was not statistically significantly changed in XAV939 treated samples (p 0.9950; Mann-Whitney U test). **F** Branching points were not statistically significantly changed in XAV939 treated samples (p 0.2306; unpaired Student’s t-test). **G** Vascular network length was not statistically significantly changed in XAV939 treated samples (p 0.1819; unpaired Student’s t-test). **H** Average vessel radius was not statistically significantly changed in XAV939 treated samples (p 0.2306; unpaired Student’s t-test). **I** Vascular complexity was not statistically significantly changed in XAV939 treated samples (p 0.9643; Mann-Whitney U test).

**Figure S12:**
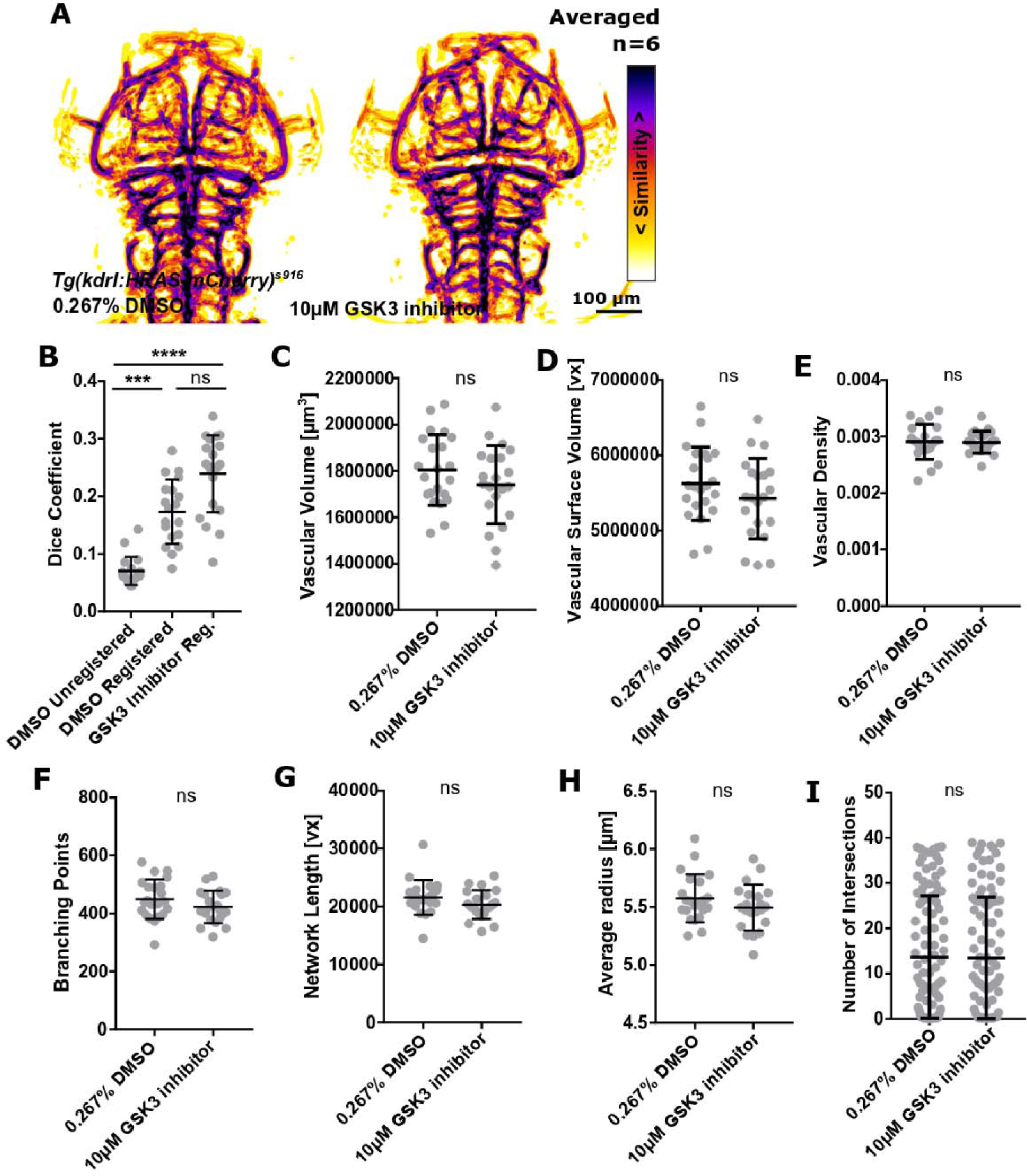
The impact of Wnt activation on vascular topology. **A** MIPs of averaged data of control and GSK3 inhibitor treated samples following segmentation and registration. **B** No statistically significant difference was found when comparing registered controls to Wnt activator treated samples (p 0.0756; control=19, GSK3 inhibitor=21; Kruskal-Wallis test). **C** Vascular volume was not statistically significantly changed in GSK3 inhibitor treated samples (p 0.2031; control=22, GSK3 inhibitor=21; unpaired t-test). **D** Vascular surface was not statistically significantly changed in GSK3 inhibitor treated samples (p 0.2108; unpaired t-test). **E** Vascular density was not statistically significantly changed in GSK3 inhibitor treated samples (p 0.8633; unpaired t-test). **F** Branching points were not statistically significantly changed in GSK3 inhibitor treated samples (p 0.1710; unpaired Student’s t-test). **G** Vascular network length was not statistically significantly changed in GSK3 inhibitor treated samples (p 0.1014; Mann-Whitney U test). **H** Average vessel radius was not statistically significantly changed in GSK3 inhibitor treated samples (p 0.2017; unpaired Student’s t-test). **I** Vascular complexity was not statistically significantly changed in GSK3 inhibitor treated samples (p 0.7764; Mann-Whitney U test).

**Figure S13:**
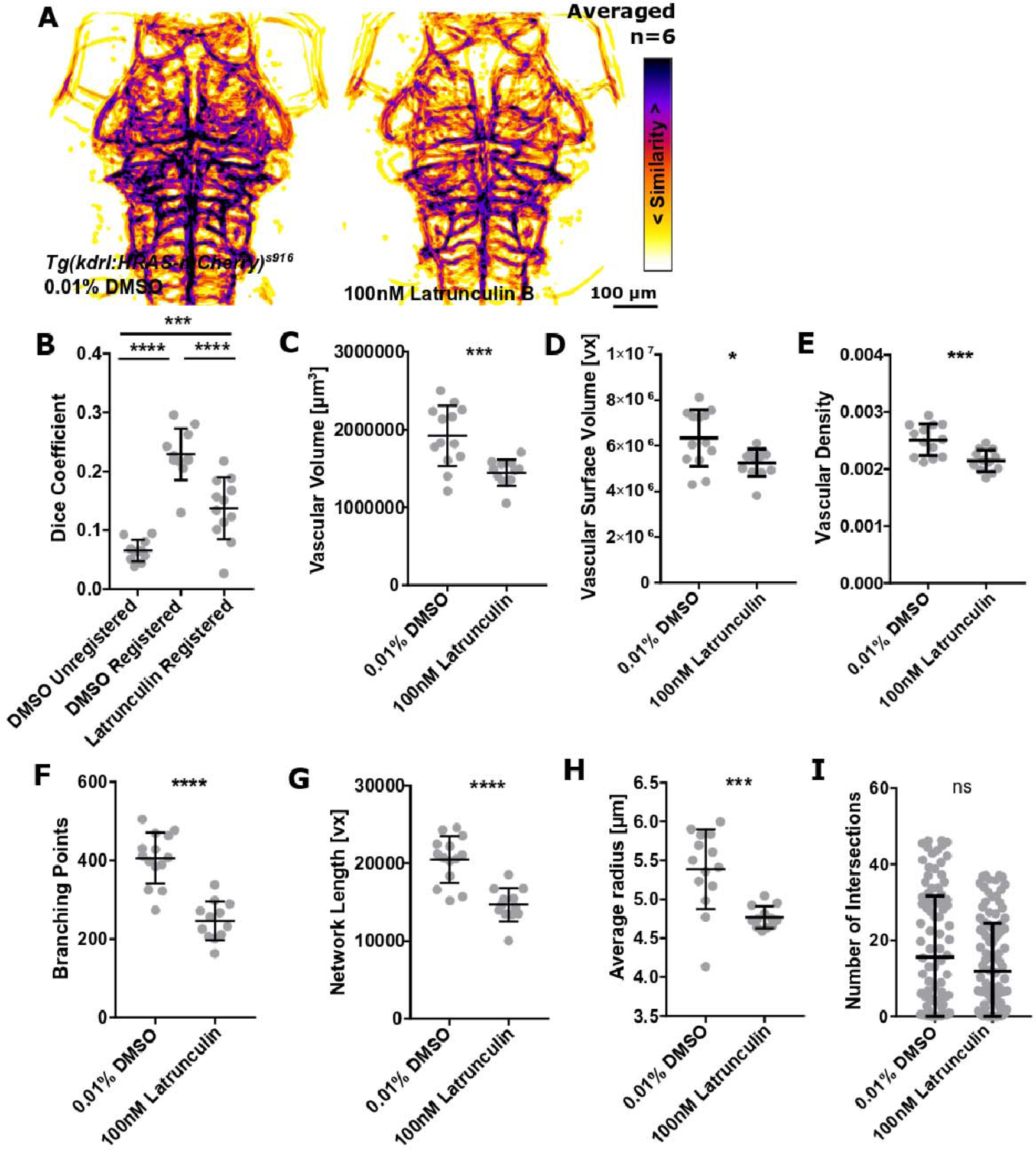
The impact of actin polymerization inhibition on vascular topology. **A** MIPs of averaged data of control and Latrunculin B treated samples following segmentation and registration. **B** A statistically significant reduction was found when comparing registered controls to F-actin inhibitor treated samples (p<0.0001; control=11, Latrunculin B=12; One-way ANOVA). **C** Vascular volume was statistically significantly decreased in Latrunculin B treated samples (p 0.0008; control=13, Latrunculin B=12; unpaired t-test). **D** Vascular surface was statistically significantly decreased in Latrunculin B treated samples (p 0.0119; unpaired t-test). **E** Vascular density was statistically significantly decreased in Latrunculin B treated samples (p 0.0008; unpaired t-test). **F** Branching points were statistically significantly decreased in Latrunculin B treated samples (p<0.0001; Mann-Whitney U test). **G** Vascular network length was statistically significantly decreased in Latrunculin B treated samples (p<0.0001; Mann-Whitney U test). **H** Average vessel radius was statistically significantly reduced in Latrunculin B treated samples (p 0.0002; Mann-Whitney U test). **I** Vascular complexity was not statistically significantly changed in Latrunculin B treated samples (p 0.1829; Mann-Whitney U test).

**Figure S14:**
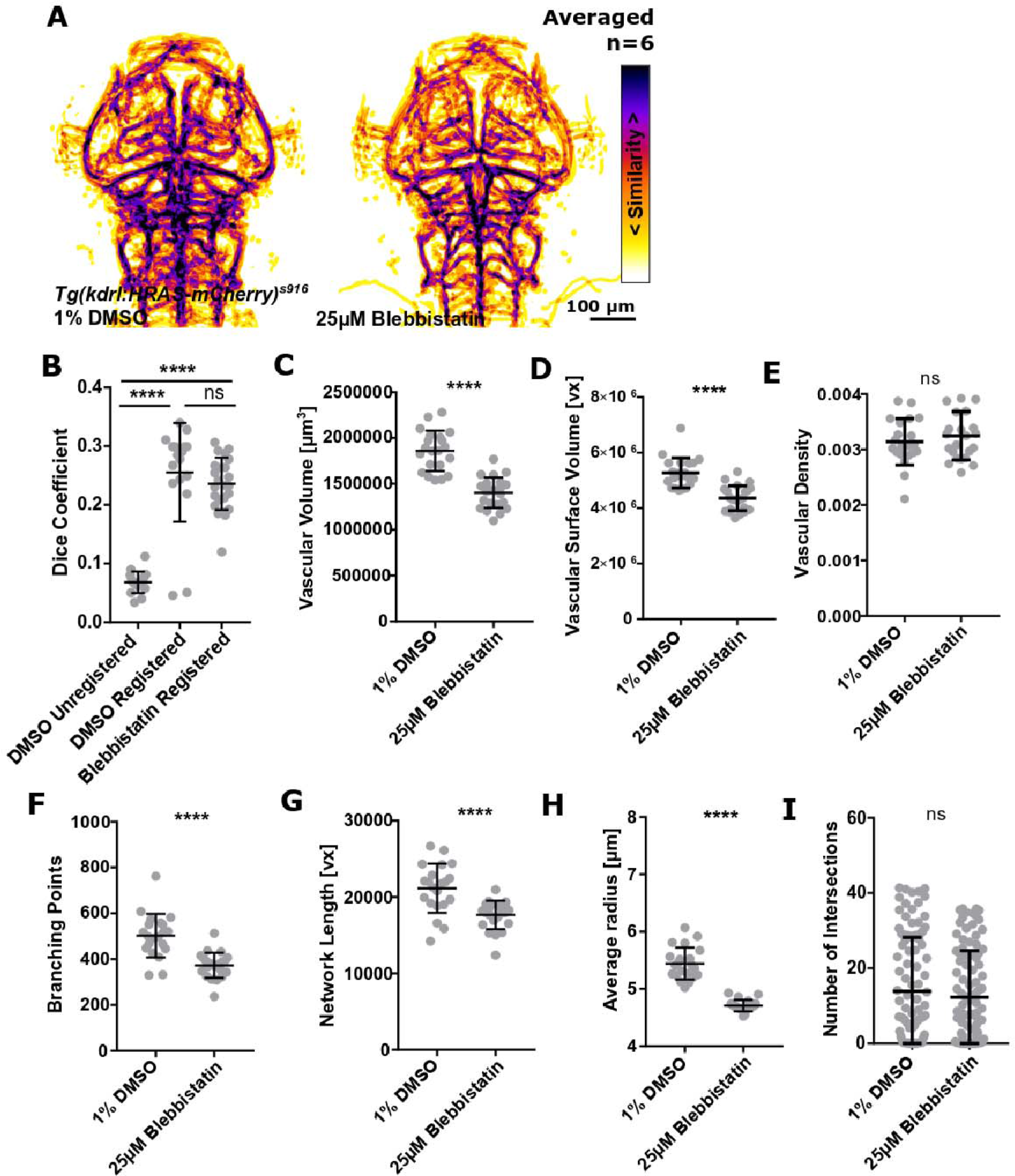
The impact of Myosin II inhibition on vascular topology. **A** MIPs of averaged data of control and Blebbistatin treated samples following segmentation and registration. **B** No statistically significant difference was found when comparing registered controls to Myosin II inhibitor treated samples (p 0.5737; control=17, Blebbistatin=23; Kruskal-Wallis test). **C** Vascular volume was statistically significantly decreased in Blebbistatin treated samples (p<0.0001; control=21, Blebbistatin=23; unpaired t-test). **D** Vascular surface was statistically significantly decreased in Blebbistatin treated samples (p<0.0001; Mann-Whitney U test). **E** Vascular density was not statistically significantly changed in Blebbistatin treated samples (p 0.4097; unpaired t-test). **F** Branching points were statistically significantly decreased in Blebbistatin treated samples (p<0.0001; unpaired Student’s t-test). **G** Vascular network length was statistically significantly decreased in Blebbistatin treated samples (p<0.0001; Mann-Whitney U test). **H** Average vessel radius was statistically significantly reduced in Blebbistatin treated samples (p <0.0001; unpaired Student’s t-test). **I** Vascular complexity was not statistically significantly changed in Blebbistatin treated samples (p 0.4909; Mann-Whitney U test).

**Figure S15:**
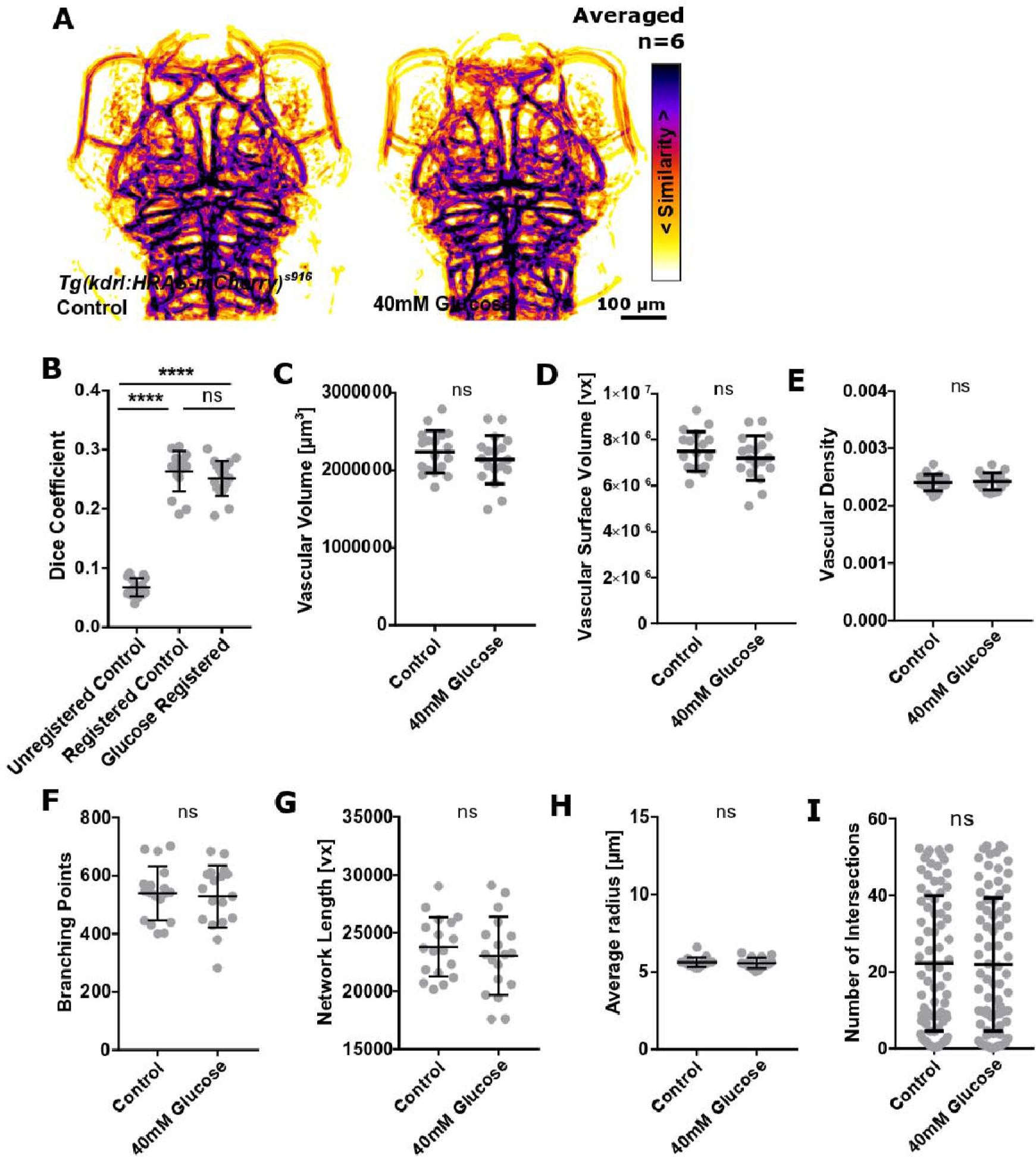
The impact of osmotic pressure increase on vascular topology. **A** MIPs of averaged data of control and glucose treated samples following segmentation and registration. **B** No statistically significant difference was found when comparing registered controls to glucose treated samples (p 0.4559; control=16, glucose=18; One-way ANOVA). **C** Vascular volume was not statistically significantly changed in glucose treated samples (p 0.3183; control=18, glucose=18; unpaired Student’s t-test). **D** Vascular surface was not statistically significantly changed in glucose treated samples (p 0.3472; unpaired t-test). **E** Vascular density was not statistically significantly changed in glucose treated samples (p 0.6678; unpaired t-test). **F** Branching points were not statistically significantly changed in glucose treated samples (p 0.7434; unpaired Student’s t-test). **G** Vascular network length was not statistically significantly changed in glucose treated samples (p 0.4488; unpaired Student’s t-test). **H** Average vessel radius was not statistically significantly changed in glucose treated samples (p 0.7016; Mann-Whitney U test). **I** Vascular complexity was not statistically significantly changed in glucose treated samples (p 0.9824; Mann-Whitney U test).

**Figure S16:**
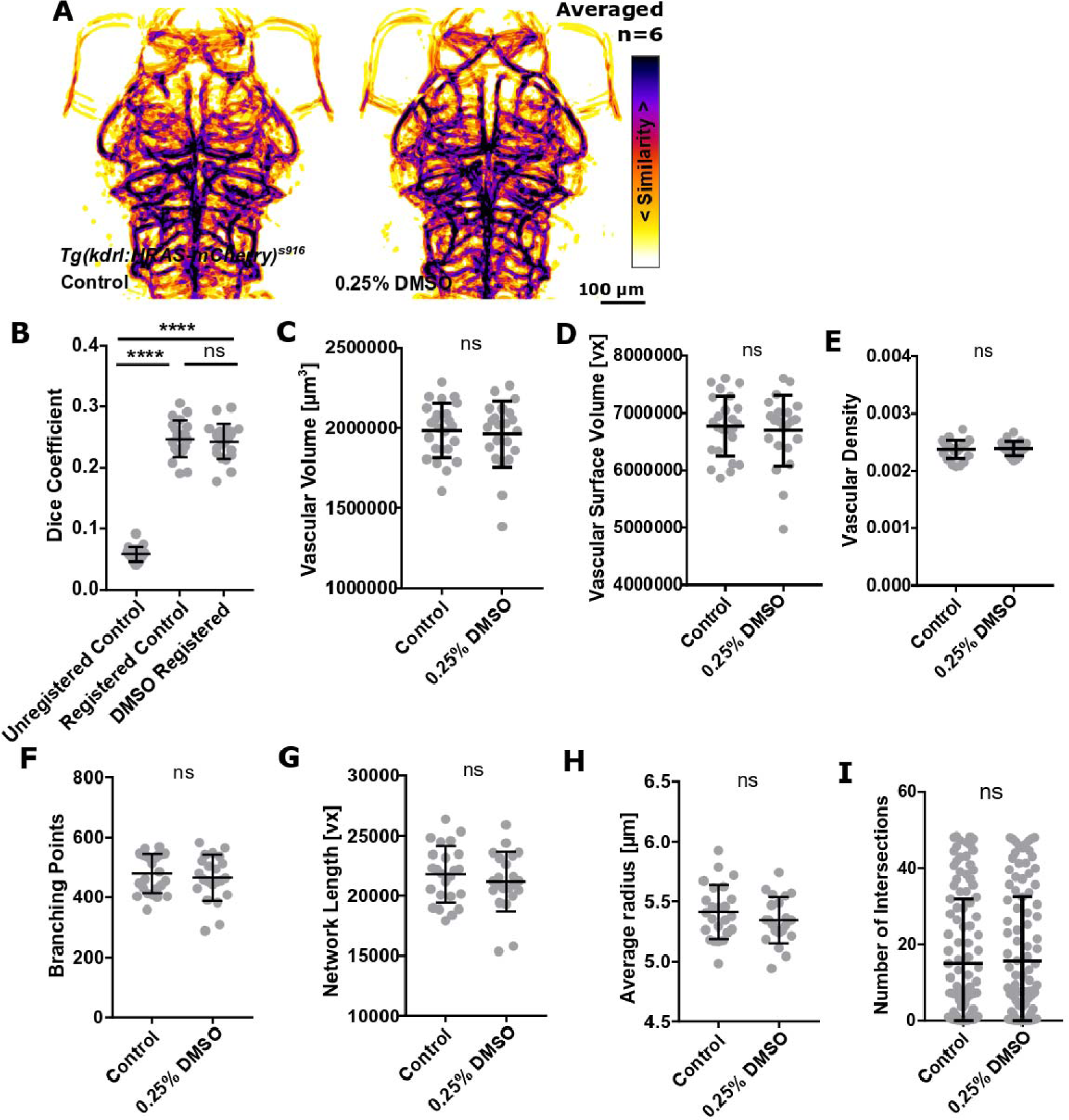
The impact of membrane rigidity decrease on vascular topology. **A** MIPs of averaged data of control and DMSO treated samples following segmentation and registration. **B** No statistically significant difference was found when comparing registered controls to DMSO treated samples (p 0.8399; control=21, DMSO=22; One-way ANOVA). **C** Vascular volume was not statistically significantly changed in DMSO treated samples (p 0.9047; control=24, DMSO=22; Mann-Whitney U test). **D** Vascular surface was not statistically significantly changed in DMSO treated samples (p>0.9999; Mann-Whitney U test). **E** Vascular density was not statistically significantly changed in DMSO treated samples (p 0.7566; unpaired t-test). **F** Branching points were not statistically significantly changed in DMSO treated samples (p 0.6909; Mann-Whitney U test). **G** Vascular network length was not statistically significantly changed in DMSO treated samples (p 0.4015; unpaired Student’s t-test). **H** Average vessel radius was not statistically significantly changed in DMSO treated samples (p 0.2676; unpaired Student’s t-test). **I** Vascular complexity was not statistically significantly changed in DMSO treated samples (p 0.5631; Mann-Whitney U test).

**Figure S17:**
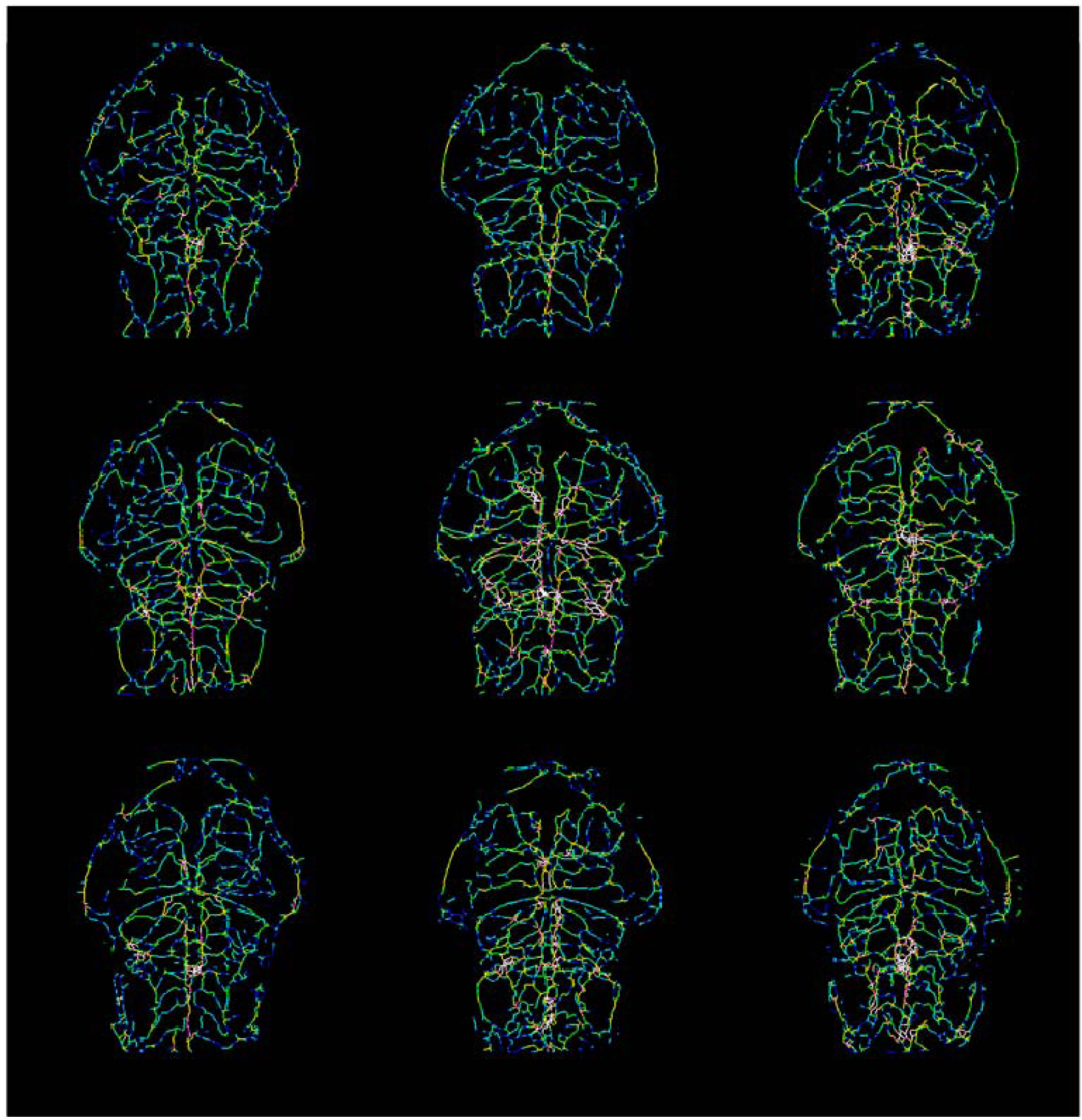
Vascular radius information is saved as one-voxel-thick representations, allowing for further examination. Image shows vessel radii on nine embryos.

**Figure S18:**
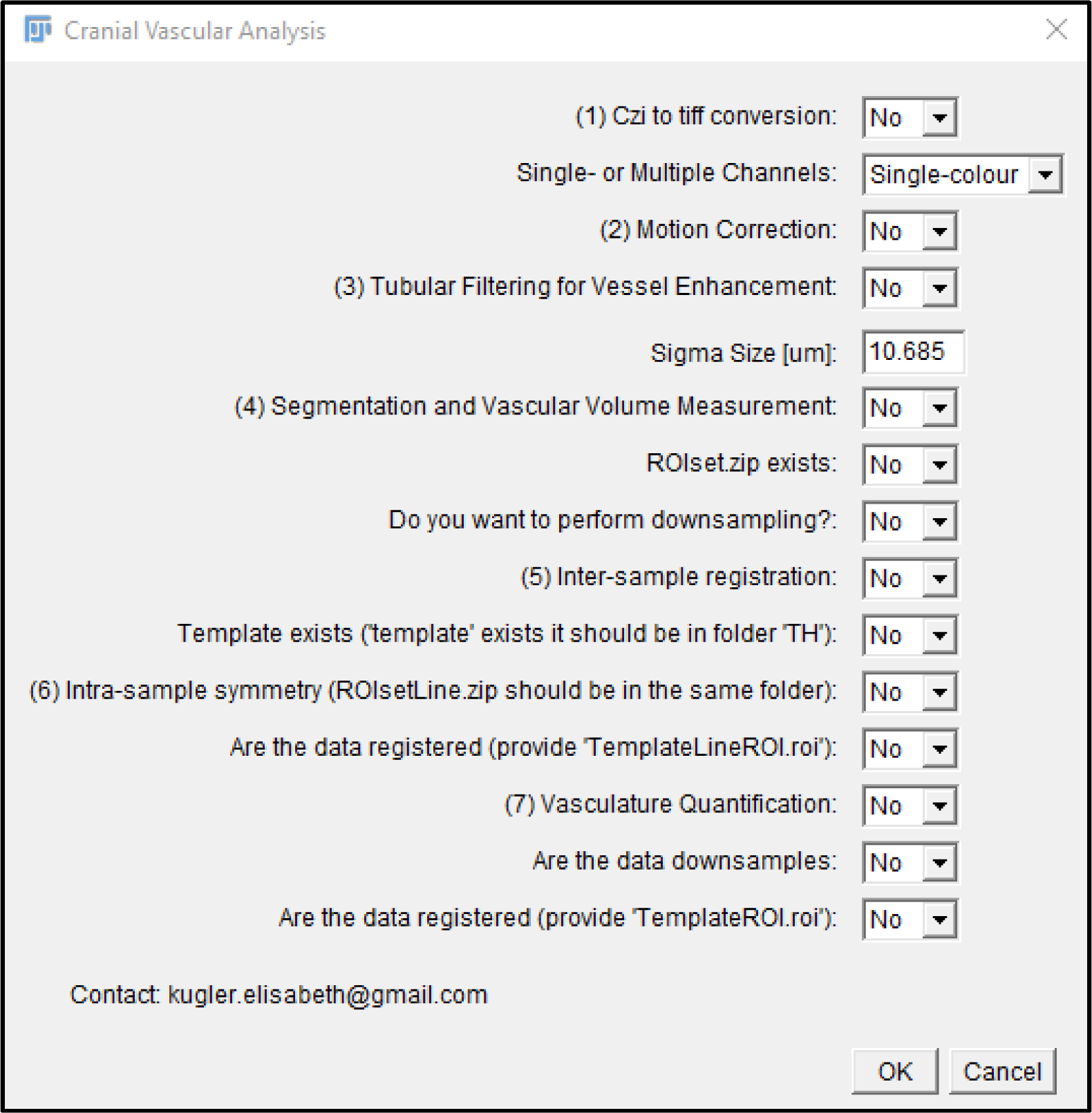
A graphical user interface (GUI) was implemented to allow dissemination and applicability of the developed workflow.

## Supplementary Videos

**Video S1 – 2dpf inter-sample registration.**

**Video S2 – 3dpf inter-sample registration.**

**Video S3 – 4dpf inter-sample registration.**

**Video S4 – 5dpf inter-sample registration.**

**Video S5 – Inter-sample registration after *tnnt2a* knock-down.** Structural similarity was statistically significantly reduced in *tnnt2a* MO (red) in comparison to control MO (white).

**Video S6 – Inter-sample registration after *jagged-1a* knock-down.** Structural similarity was not statistically significantly altered in *jagged-1a* MO (red) in comparison to control MO (white).

**Video S7 – Inter-sample registration after *jagged-1b* knock-down.** Structural similarity was not statistically significantly altered in *jagged-1b* MO (red) in comparison to control MO (white).

**Video S8 – Inter-sample registration after *dll4* knock-down.** Structural similarity was statistically significantly reduced in *dll4* MO (red) in comparison to control MO (white).

**Video S9 – Inter-sample registration after *notch1b* knock-down.** Structural similarity was not statistically significantly altered in *notch1b* MO (red) in comparison to control MO (white).

**Video S10 – Inter-sample registration after *ccbe1* knock-down.** Structural similarity was not statistically significantly altered in *ccbe1* MO (red) in comparison to control MO (white).

**Video S11 – Inter-sample registration after VEGF inhibition.** Structural similarity was not statistically significantly altered upon AV951 treatment (red) in comparison to controls (white).

**Video S12 – Inter-sample registration after Notch inhibition.** Structural similarity was not statistically significantly altered upon DAPT treatment (red) in comparison to controls (white).

**Video S13 – Inter-sample registration after NOS inhibition.** Structural similarity was not statistically significantly altered upon L-NAME treatment (red) in comparison to controls (white).

**Video S14 – Inter-sample registration after Wnt inhibition.** Structural similarity was not statistically significantly altered upon XAV939 treatment (red) in comparison to controls (white).

**Video S15 – Inter-sample registration after Wnt activation.** Structural similarity was not statistically significantly altered upon GSK3 inhibitor treatment (red) in comparison to controls (white).

**Video S16 – Inter-sample registration after actin polymerization inhibition.** Structural similarity was statistically reduced upon Latrunculin B treatment (red) in comparison to controls (white).

**Video S17 – Inter-sample registration after Myosin II inhibition.** Structural similarity was not statistically significantly altered upon Blebbistatin treatment (red) in comparison to controls (white).

**Video S18 – Inter-sample registration after osmotic pressure increase.** Structural similarity was not statistically significantly altered upon glucose treatment (red) in comparison to controls (white).

**Video S19 – Inter-sample registration after membrane rigidity increase.** Structural similarity was not statistically significantly altered upon DMSO treatment (red) in comparison to controls (white).

**Video S20 – 3dpf intra-sample left-right symmetry.** 3D rendering of overlapped left (green) and right (magenta) vasculature at 3dpf.

## References

1. Chávez, M. N., Aedo, G., Fierro, F. A., Allende, M. L. & Egaña, J. T. Zebrafish as an Emerging Model Organism to Study Angiogenesis in Development and Regeneration. Front Physiol 7, (2016).

2. Chico, T. J. A., Ingham, P. W. & Crossman, D. C. Modeling Cardiovascular Disease in the Zebrafish. Trends in Cardiovascular Medicine 18, 150–155 (2008).

3. Huisken, J., Swoger, J., Del Bene, F., Wittbrodt, J. & Stelzer, E. H. K. Optical sectioning deep inside live embryos by selective plane illumination microscopy. Science 305, 1007–1009 (2004).

4. Gut, P., Reischauer, S., Stainier, D. Y. R. & Arnaout, R. Little Fish, Big Data: Zebrafish as a Model for Cardiovascular and Metabolic Disease. Physiological Reviews 97, 889–938 (2017).

5. Haixiang, G., et al. Learning from class-imbalanced data: Review of methods and applications. Expert Systems with Applications 73, 220–239 (2017).

6. Rahman, M. M. & Davis, D. N. Addressing the Class Imbalance Problem in Medical Datasets. IJMLC 224–228 (2013) doi:10.7763/IJMLC.2013.V3.307.

7. Tetteh, G., et al. DeepVesselNet: Vessel Segmentation, Centerline Prediction, and Bifurcation Detection in 3-D Angiographic Volumes. arXiv:1803.09340 [cs] (2019).

8. Tam, S. J. et al. Death Receptors DR6 and TROY Regulate Brain Vascular Development. Developmental Cell 22, 403–417 (2012).

9. Chen, Q., et al. Haemodynamics-Driven Developmental Pruning of Brain Vasculature in Zebrafish. PLOS Biology (2012).

10. Kugler, E., Chico, T. & Armitage, P. Image Analysis in Light Sheet Fluorescence Microscopy Images of Transgenic Zebrafish Vascular Development. in Nixon M., Mahmoodi S., Zwiggelaar R. (eds) Medical Image Understanding and Analysis. MIUA 2018. vol. Communications in Computer and Information Science, vol 894 343–353 (Springer, Cham, 2018).

11. Kugler, E., Plant, K., Chico, T. & Armitage, P. Enhancement and Segmentation Workflow for the Developing Zebrafish Vasculature. Journal of Imaging 5, 14 (2019).

12. Daetwyler, S., Günther, U., Modes, C. D., Harrington, K. & Huisken, J. Multi-sample SPIM image acquisition, processing and analysis of vascular growth in zebrafish. Development dev.173757 (2019) doi:10.1242/dev.173757.

13. Kugler, E., Chico, T. & Armitage, P. A. Validating Segmentation of the Zebrafish Vasculature. in Medical Image Understanding and Analysis (eds. Zheng, Y., Williams, B. M. & Chen, K.) 270–281 (Springer International Publishing, 2020). doi:10.1007/978-3-030-39343-4_23.

14. Kugler, E. C., Rampun, A., Chico, T. & Armitage, P. Segmentation of the Zebrafish Brain Vasculature from Light Sheet Fluorescence Microscopy Datasets. http://biorxiv.org/lookup/doi/10.1101/2020.07.21.213843 (2020) doi:10.1101/2020.07.21.213843.

15. Sehnert, A. J. et al. Cardiac troponin T is essential in sarcomere assembly and cardiac contractility. Nat. Genet. 31, 106–110 (2002).

16. Geudens, I. et al. Role of delta-like-4/Notch in the formation and wiring of the lymphatic network in zebrafish. Arterioscler. Thromb. Vasc. Biol. 30, 1695–1702 (2010).

17. Quillien, A. et al. Distinct Notch signaling outputs pattern the developing arterial system. Development 141, 1544–1552 (2014).

18. Guen, L. L. et al. Ccbe1 regulates Vegfc-mediated induction of Vegfr3 signaling during embryonic lymphangiogenesis. Development 141, 1239–1249 (2014).

19. Nakamura, K. et al. KRN951, a highly potent inhibitor of vascular endothelial growth factor receptor tyrosine kinases, has antitumor activities and affects functional vascular properties. Cancer Res. 66, 9134–9142 (2006).

20. Geling, A., Steiner, H., Willem, M., Bally-Cuif, L. & Haass, C. A gamma-secretase inhibitor blocks Notch signaling in vivo and causes a severe neurogenic phenotype in zebrafish. EMBO Rep 3, 688–694 (2002).

21. Ziche, M. et al. Nitric oxide synthase lies downstream from vascular endothelial growth factor-induced but not basic fibroblast growth factor-induced angiogenesis. J. Clin. Invest. 99, 2625–2634 (1997).

22. Huang, S.-M. A. et al. Tankyrase inhibition stabilizes axin and antagonizes Wnt signalling. Nature 461, 614–620 (2009).

23. Du, J. et al. A kinesin signaling complex mediates the ability of GSK-3beta to affect mood- associated behaviors. Proc Natl Acad Sci U S A 107, 11573–11578 (2010).

24. Morton, W. M., Ayscough, K. R. & McLaughlin, P. J. Latrunculin alters the actin-monomer subunit interface to prevent polymerization. Nature Cell Biology 2, 376–378 (2000).

25. Kovács, M., Tóth, J., Hetényi, C., Málnási-Csizmadia, A. & Sellers, J. R. Mechanism of blebbistatin inhibition of myosin II. J. Biol. Chem. 279, 35557–35563 (2004).

26. Schindelin, J. et al. Fiji - an Open Source platform for biological image analysis. Nat Methods 9, (2012).

27. Chi, N. C. et al. Foxn4 directly regulates tbx2b expression and atrioventricular canal formation. Genes Dev 22, 734–739 (2008).

28. Sato, Y. et al. 3D multi-scale line filter for segmentation and visualization of curvilinear structures in medical images. in CVRMed-MRCAS’97 213–222 (Springer, Berlin, Heidelberg, 1997). doi:10.1007/BFb0029240.

29. Otsu, N. A threshold selection method from gray-level histograms. Trans. Sys.Man. 9, 62–66 (1979).

30. Schmid, B. Computational tools for the segmentation and registration of confocal brain images of Drosophila melanogaster. (Bayerischen Julius-Maximilians-Universitaet Wuerzburg, 2010).

31. Jenett, A., Schindelin, J. E. & Heisenberg, M. The Virtual Insect Brain protocol: creating and comparing standardized neuroanatomy. BMC Bioinformatics 7, 544 (2006).

32. Leslie, J. D. et al. Endothelial signalling by the Notch ligand Delta-like 4 restricts angiogenesis. Development 134, 839–844 (2007).

33. Lobov, I. B. et al. Delta-like ligand 4 (Dll4) is induced by VEGF as a negative regulator of angiogenic sprouting. PNAS 104, 3219–3224 (2007).

34. Hogan, B. M. et al. Ccbe1 is required for embryonic lymphangiogenesis and venous sprouting. Nat. Genet. 41, 396–398 (2009).

35. Bonet, F. et al. CCBE1 is required for coronary vessel development and proper coronary artery stem formation in the mouse heart. Developmental Dynamics 247, 1135–1145 (2018).

36. Bower, N. I. et al. Vegfd modulates both angiogenesis and lymphangiogenesis during zebrafish embryonic development. Development 144, 507–518 (2017).

37. Ellis, L. M. & Hicklin, D. J. VEGF-targeted therapy: mechanisms of anti-tumour activity. Nature Reviews Cancer 8, 579–591 (2008).

38. Gerhardt, H. et al. VEGF guides angiogenic sprouting utilizing endothelial tip cell filopodia. J Cell Biol 161, 1163–1177 (2003).

39. Bray, S. J. Notch signalling: a simple pathway becomes complex. Nat. Rev. Mol. Cell Biol. 7, 678–689 (2006).

40. Watson, O. et al. Blood flow suppresses vascular Notch signalling via dll4 and is required for angiogenesis in response to hypoxic signalling. Cardiovasc Res 100, 252–261 (2013).

41. Fritsche, R., Schwerte, T. & Pelster, B. Nitric oxide and vascular reactivity in developing zebrafish, Danio rerio. Am. J. Physiol. Regul. Integr. Comp. Physiol. 279, R2200–2207 (2000).

42. Stenman, J. M. et al. Canonical Wnt Signaling Regulates Organ-Specific Assembly and Differentiation of CNS Vasculature. Science 322, 1247–1250 (2008).

43. Parmalee, N. L. & Kitajewski, J. Wnt Signaling in Angiogenesis. Curr Drug Targets 9, 558–564 (2008).

44. Hübner, K. et al. Wnt/beta-catenin signaling regulates VE-cadherin-mediated anastomosis of brain capillaries by counteracting S1pr1 signaling. Nature Communications 9, 4860 (2018).

45. Sacharidou, A., Stratman, A. N. & Davis, G. E. Molecular Mechanisms Controlling Vascular Lumen Formation in Three-Dimensional Extracellular Matrices. CTO 195, 122–143 (2012).

46. Scallan, J., Huxley, V. H. & Korthuis, R. J. Pathophysiology of Edema Formation. Capillary Fluid Exchange: Regulation, Functions, and Pathology (Morgan & Claypool Life Sciences, 2010).

47. Severinghaus, J. W. Hypothetical roles of angiogenesis, osmotic swelling, and ischemia in high- altitude cerebral edema. J. Appl. Physiol. 79, 375–379 (1995).

48. Chhabria, K. et al. The effect of hyperglycemia on neurovascular coupling and cerebrovascular patterning in zebrafish. J. Cereb. Blood Flow Metab. 271678X18810615 (2018) doi:10.1177/0271678X18810615.

49. de Ménorval, M.-A., Mir, L. M., Fernández, M. L. & Reigada, R. Effects of dimethyl sulfoxide in cholesterol-containing lipid membranes: a comparative study of experiments in silico and with cells. PLoS ONE 7, e41733 (2012).

50. Glass, C. A., Perrin, R. M., Pocock, T. M. & Bates, D. O. Transient osmotic absorption of fluid in microvessels exposed to low concentrations of dimethyl sulfoxide. Microcirculation 13, 29–40 (2006).

51. Randlett, O. et al. Whole-brain activity mapping onto a zebrafish brain atlas. Nat. Methods 12, 1039–1046 (2015).

52. Marquart, G. D. et al. High-precision registration between zebrafish brain atlases using symmetric diffeomorphic normalization. Gigascience 6, 1–15 (2017).

53. Meyer, A. & Schartl, M. Gene and genome duplications in vertebrates: the one-to-four (-to-eight in fish) rule and the evolution of novel gene functions. Curr. Opin. Cell Biol. 11, 699–704 (1999).

54. Phng, L.-K. & Gerhardt, H. Angiogenesis: A Team Effort Coordinated by Notch. Developmental Cell 16, 196–208 (2009).

55. Hogan, B. M. & Schulte-Merker, S. How to Plumb a Pisces: Understanding Vascular Development and Disease Using Zebrafish Embryos. Developmental Cell 42, 567–583 (2017).

56. Dreosti, E., Vendrell Llopis, N., Carl, M., Yaksi, E. & Wilson, S. W. Left-Right Asymmetry Is Required for the Habenulae to Respond to Both Visual and Olfactory Stimuli. Current Biology 24, 440–445 (2014).

57. Westerfield, M. The Zebrafish Book: A Guide for Laboratory use of Zebrafish (Brachydanio rerio). (University of Oregon Press, 1993).

58. Kugler, E. C. et al. Cerebrovascular endothelial cells form transient Notch-dependent cystic structures in zebrafish. EMBO reports 20, e47047 (2019).

59. Lowe, D. Object Recognition from Local Scale-Invariant Features. in Proc. of the International Conference on Computer Vision (1999).

60. Lowe, D. G. Distinctive Image Features from Scale-Invariant Keypoints. International Journal of Computer Vision 60, 91–110 (2004).

61. Legland, D., Arganda-Carreras, I. & Andrey, P. MorphoLibJ: integrated library and plugins for mathematical morphology with ImageJ. Bioinformatics 32, 3532–3534 (2016).

62. Canny, J. A computational Approach to Edge Detection. IEEE Transactions on Pattern Analysis and Machine Intelligence PAMI-8, 679–698 (1986).

63. Borgefors, G. On Digital Distance Transforms in Three Dimensions. Computer Vision and Image Understanding 64, 368–376 (1996).

64. Lee, T. C., Kashyap, R. L. & Chu, C. N. Building Skeleton Models via 3-D Medial Surface Axis Thinning Algorithms. CVGIP: Graphical Models and Image Processing 56, 462–478 (1994).

65. Arganda-Carreras, I., Fernández-González, R., Muñoz-Barrutia, A. & Ortiz-De-Solorzano, C. 3D reconstruction of histological sections: Application to mammary gland tissue. Microsc. Res. Tech. 73, 1019–1029 (2010).

66. Sholl, D. A. Dendritic organization in the neurons of the visual and motor cortices of the cat. J. Anat. 87, 387–406 (1953).

67. Bird, A. D. & Cuntz, H. Dissecting Sholl Analysis into Its Functional Components. Cell Reports 27, 3081–3096.e5 (2019).

68. Ferreira, T. A. et al. Neuronal morphometry directly from bitmap images. Nature Methods 11, 982–984 (2014).

69. Metsalu, T. & Vilo, J. ClustVis: a web tool for visualizing clustering of multivariate data using Principal Component Analysis and heatmap. Nucleic Acids Res 43, W566–W570 (2015).

